# Epigenomic and functional characterization of a core DNA methyltransferase in the human pathogen *Clostridium difficile*

**DOI:** 10.1101/398891

**Authors:** Pedro H. Oliveira, John W. Ribis, Elizabeth M. Garrett, Dominika Trzilova, Alex Kim, Ognjen Sekulovic, Edward A. Mead, Theodore Pak, Shijia Zhu, Gintaras Deikus, Marie Touchon, Colleen Beckford, Nathalie E. Zeitouni, Deena Altman, Elizabeth Webster, Irina Oussenko, Supinda Bunyavanich, Aneel K. Aggarwal, Ali Bashir, Gopi Patel, Camille Hamula, Shirish Huprikar, Eric E. Schadt, Robert Sebra, Harm van Bakel, Andrew Kasarskis, Rita Tamayo, Aimee Shen, Gang Fang

**Affiliations:** Department of Genetics and Genomic Sciences, Institute for Genomics and Multiscale Biology, Mount Sinai School of Medicine, New York, New York, United States of America; Department of Molecular Biology and Microbiology, Tufts University School of Medicine, Boston, Massachusetts, United States of America; Department of Microbiology and Immunology, University of North Carolina at Chapel Hill School of Medicine, Chapel Hill, North Carolina, USA; Microbial Evolutionary Genomics, Institut Pasteur, 25–28 rue du Docteur Roux, Paris, 75015, France; CNRS, UMR3525, 25–28 rue du Docteur Roux, Paris, 75015, France; Department of Medicine, Infectious Disease, Mount Sinai School of Medicine, New York, New York, United States of America; Department of Pharmacological Sciences and Department of Oncological Sciences, Mount Sinai School of Medicine, New York, New York, United States of America

**Author notes:** Email: Aimee Shen; Gang Fang.

**Keywords:** DNA methylation, SMRT sequencing, sporulation, restriction-modification systems

## Abstract

*Clostridioides difficile* is a leading cause of health care-associated infections. Although significant progress has been made in the understanding of its genome, the epigenome of *C. difficile* and its functional impact has not been systematically explored. Here, we performed the first comprehensive DNA methylome analysis of *C. difficile* using 36 human isolates and observed great epigenomic diversity. We discovered an orphan DNA methyltransferase with a well-defined specificity whose corresponding gene is highly conserved across our dataset and in all ~300 global *C. difficile* genomes examined. Inactivation of the methyltransferase gene negatively impacted sporulation, a key step in *C. difficile* disease transmission, consistently supported by multi-omics data, genetic experiments, and a mouse colonization model. Further experimental and transcriptomic analysis also suggested that epigenetic regulation is associated with cell length, biofilm formation, and host colonization. These findings open up a new epigenetic dimension to characterize medically relevant biological processes in this critical pathogen. This work also provides a set of methods for comparative epigenomics and integrative analysis, which we expect to be broadly applicable to bacterial epigenomics studies.

## Introduction

*Clostridioides* (formerly *Clostridium*) *difficile* is a spore-forming Gram-positive obligate anaerobe and the leading cause of nosocomial antibiotic-associated disease in the developed world^1^. Clinical symptoms of *C. difficile* infection (CDI) in humans range in severity from mild self-limiting diarrhea to severe, life-threatening inflammatory conditions, such as pseudomembranous colitis or toxic megacolon. Since the vegetative form of *C. difficile* cannot survive in the presence of oxygen, the bacterium is transmitted via the fecal/oral route as a metabolically dormant spore^2^. In the intestinal environment, these spores subsequently germinate into actively growing, toxin-producing vegetative cells that are responsible for disease pathology^3^. CDI progresses in an environment of host microbiota dysbiosis, which disrupts the colonization resistance typically provided by a diverse intestinal microbiota^4^. In the last two decades, there has been a dramatic rise in outbreaks with increased mortality and morbidity due in part to the emergence of epidemic-associated strains with enhanced growth^5,6^, toxin production^7^, and antibiotic resistance^8^*. C. difficile* was responsible for half a million infections in the United States in 2011, with 29,000 individuals dying within 30 days of the initial diagnosis^9^. Those most at risk are older adults, particularly those who take antibiotics that perturb the normally protective intestinal microbiota.

Despite the significant progress achieved in the understanding of *C. difficile* physiology, genetics, and genomic evolution^10,11^, the roles played by epigenetic factors, namely DNA methylation, have not been systematically studied^12,13^. In the bacterial kingdom, there are three major forms of DNA methylation: N6-methyladenine (6mA, the most prevalent form representing ~80%), N4-methylcytosine (4mC), and 5-methylcytosine (5mC). Although bacterial DNA methylation is most commonly associated with restriction-modification (R-M) systems that defend hosts against invading foreign DNA^14,15^, increasing evidence suggests that DNA methylation also regulates a number of biological processes such as DNA replication and repair, cell cycle, chromosome segregation and gene expression, among others^16–21^. Efficient high-resolution mapping of bacterial DNA methylation events has only recently become possible with the advent of Single Molecule Real-Time sequencing (SMRT-seq)^22,23^, which can detect all three types of DNA methylation, albeit at different signal-to-noise ratios: high for 6mA, medium for 4mC, and low for 5mC. This technique enabled the characterization of the first bacterial methylomes^24,25^, and since then, more than 2,100 (as of 06/2019) have been mapped, heralding a new era of “bacterial epigenomics”^26^.

Herein, we mapped and characterized DNA methylomes of 36 human *C. difficile* isolates using SMRT-seq and comparative epigenomics. We observed great epigenomic diversity across *C. difficile* isolates, as well as the presence of a highly conserved methyltransferase (MTase). Inactivation of this MTase resulted in a functional impact on sporulation, a key step in *C. difficile* transmission, consistently supported by multi-omics data and genetic experiments. Further experimental and integrative transcriptomic analysis suggested that epigenetic regulation by DNA methylation also modulates *C. difficile* cell length, host colonization and biofilm formation. Finally, the epigenomic landscape of *C. difficile* also allowed us to perform a data-driven joint analysis of multiple defense systems and their contribution to gene flux. These discoveries are expected to stimulate future investigations along a new epigenetic dimension to characterize and potentially repress medically relevant biological processes in this critical pathogen.

## Results

### Methylome analysis reveals great epigenomic diversity in *C. difficile*

From an ongoing Pathogen Surveillance Project at Mount Sinai Medical Center, 36 *C. difficile* isolates were collected from fecal samples of infected patients (Supplementary Table 1). A total of 15 different MLST sequence types (STs) belonging to clades 1 (human and animal, HA1) and 2 (so-called hypervirulent or epidemic)^27^ are represented in our dataset (Fig. 1a). Using SMRT-seq with long library size selection, *de novo* genome assembly was achieved at high quality (Supplementary Table 1). Methylation motifs were found using the SMRTportal protocol (Materials and Methods). We found a total of 17 unique high-quality methylation motifs in the 36 genomes (average of 2.6 motifs per genome) (Fig. 1a, Supplementary Table 2a). The large majority of target motifs were of 6mA type, one motif (TAA<u>C</u>TG) belonged to the 4mC type, and no confident 5mC motifs were detected (Supplementary Text). Like most bacterial methylomes, >95% of the 6mA and 4mC motif sites were methylated (Fig. 1b, Supplementary Table 2a).

**Fig. 1.**
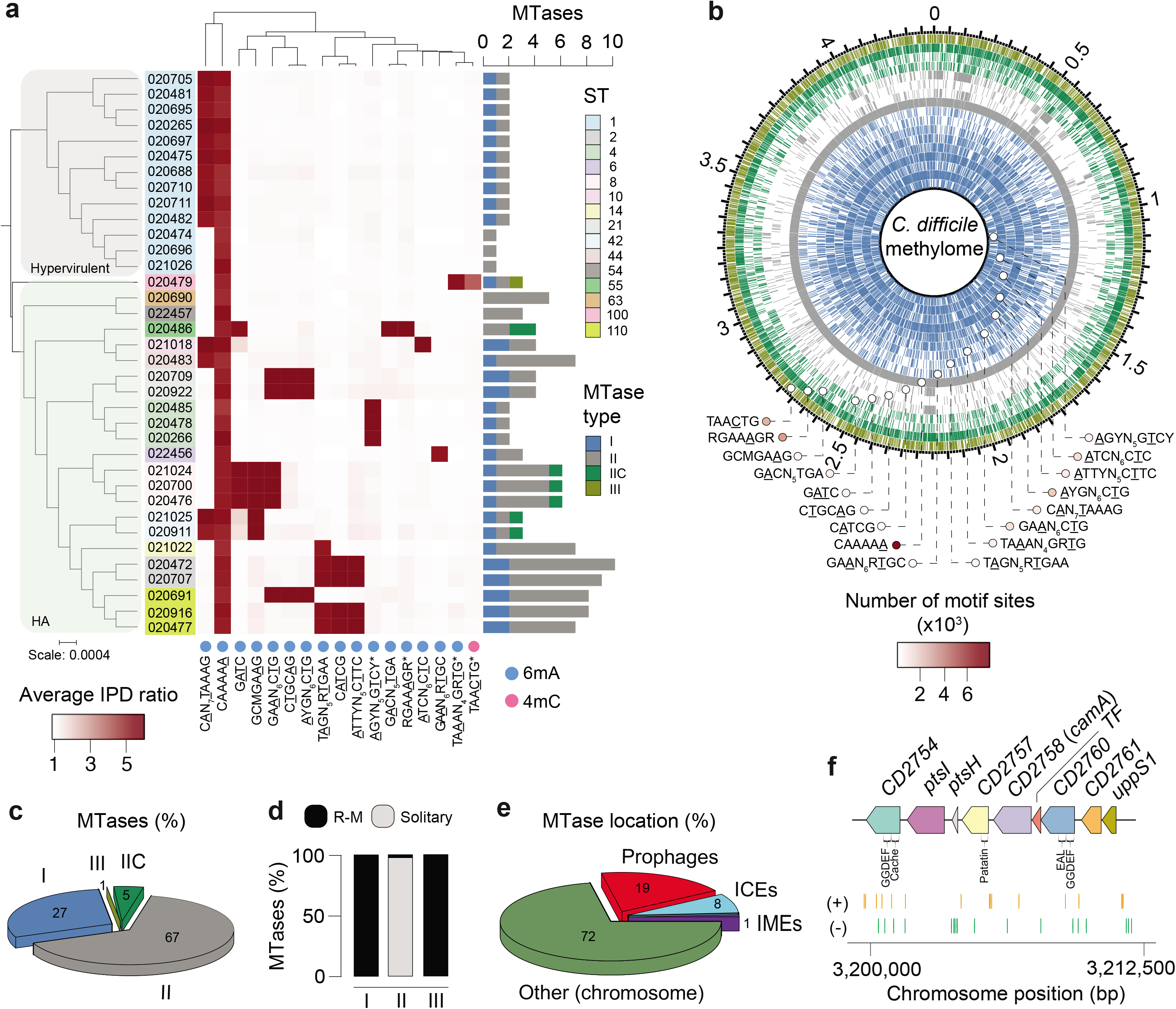
Methylomes of the 36 *C. difficile* strains. (a) Phylogenetic tree of the 36 *C. difficile* strains colored by clade (hypervirulent, human and animal (HA) associated) and MLST sequence type (ST). Heatmap depicting the landscape of methylated motifs per genome, and their average interpulse duration (IPD) ratio. Asterisks refer to new motifs not previously listed in the reference database REBASE. Methylated bases are underlined. The CAAAA<u>A</u> motif was consistently methylated across isolates. Barplot indicates the number and types of active MTases detected per genome. In Type IIC systems, MTase and REase are encoded in the same polypeptide. (b) Representation of the *C. difficile* methylome. Shown are the positions of all methylation motif sites in the reference genome of *C. difficile* 630, colored according to MTase type. Also shown are the average motif occurrences per genome (across the 36 isolates). (c) % of MTases detected according to type. (d) % MTases pertaining to complete R-M systems or without cognate REase (solitary). (e) Breakdown of MTases by location: Integrative Mobile Elements (IMEs), Integrative Conjugative Elements (ICEs), prophages, and other (within the chromosome). No hits were obtained in plasmids. (f) Immediate genomic context of *camA*. The example shown (including coordinates) refers to the reference genome of *C. difficile* 630. + / − signs correspond to the sense and antisense strands respectively. Vertical bars correspond to the distribution of the CAAAA<u>A</u> motif. *CD2754*: phosphodiesterase with a GGDEF domain (PF00990) and a cache domain (PF02743); *ptsI* and *ptsH* belong to a phosphotransferase (PTS) system; *CD2757*: patatin-like phospholipase (PF01734); *CD2758* (*camA*): Type II MTase; *CD2759*: Rrf2-type transcriptional regulator; *CD2760*: phosphodiesterase with a GGDEF domain (PF00990) and a conserved EAL domain (PF00563)^87^; *CD2761*: N-acetylmuramoyl-L-alanine amidase; *CD2762*: undecaprenyl diphosphate synthase.

Genomes pertaining to the same ST tend to have more similar sets of methylation motifs relative to those from different STs. Those belonging to ST-2, ST-8, ST-21, and ST-110 showed the highest motif diversities. One 6mA motif, CAAAA<u>A</u>, was present across all genomes, which led us to hypothesize that 6mA methylation events at this motif, and its corresponding MTase, play an important and conserved function in *C. difficile*.

### A DNA methyltransferase and its target motif are ubiquitous in *C. difficile*

Motivated by the consistent presence of the methylation motif CAAAA<u>A</u> across all the *C. difficile* isolates, we proceeded to examine the encoded MTases. From the 36 genome assemblies, we identified a total of 139 MTase genes (average of 3.9 per genome) (Fig. 1a, Supplementary Table 2b) representing all the four major types^28^. 38 MTases (27%) belong to Type I R-M systems, and 100 MTases (72%) belong to Type II (Fig. 1c). All but one of the Type II MTases are solitary, *i.e*., devoid of a cognate restriction endonuclease (REase) (Fig. 1d). The least abundant MTases (1%) belong to Type III R-M systems (Fig. 1c). A Type IV (no cognate MTase) McrBC REase gene was also present in all genomes (Supplementary Table 2b). 28% of MTase genes were located in mobile genetic elements (MGEs) (19% in prophages and 9% in integrative conjugative/mobile elements), while the large majority was encoded in other chromosomal regions (Fig. 1e, Supplementary Tables 2b-d). No MTase genes were found in plasmids. We further found multiple additional defense systems (*e.g*., abortive infection systems, CRISPR-Cas, toxin-antitoxin), and performed an integrative analysis with R-M systems in relation to host defense and gene flux (Supplementary Figs. 1a-e, Supplementary Fig. 2, Supplementary Tables 3a-f, Supplementary Text), such as that involving phages (Supplementary Fig. 3, Supplementary Table 3g, Supplementary Text).

Consistent with the presence of a highly conserved CAAAA<u>A</u> motif, we identified a Type II 6mA solitary DNA MTase (577 aa) present across isolates (Fig. 1f, Supplementary Table 2b). This MTase is encoded by *CD2758* in *C. difficile* 630, a reference strain that was isolated from a *C. difficile* outbreak in Switzerland^10,29^. Here we have named CD2758 as CamA (***<u>C</u>**. **<u>d</u>**ifficile* **<u>a</u>**denine **<u>m</u>**ethyltransferase **<u>A</u>**). The ubiquity of this MTase was not restricted to the 36 isolates, as we were able to retrieve orthologs in a list of ~300 global *C. difficile* isolates from GenBank (Supplementary Table 4). REBASE also showed functional orthologs of *camA* only in very few other *Clostridiales* and *Fusobacteriales* (Supplementary Figs. 4a, b), suggesting that this MTase is fairly unique to *C. difficile*. The genomic context of *camA* is largely conserved across strains (Supplementary Fig. 4c), located ~25 kb upstream of the S-layer biogenesis locus (Supplementary Fig. 4d). Several of the genes flanking *camA* (including itself) are part of the *C. difficile* core-genome (see below), suggesting that they may play biological roles fundamental to *C. difficile*. Hence, CamA is a solitary Type II 6mA MTase that specifically recognizes the CAAAA<u>A</u> motif and is ubiquitous and fairly unique to *C. difficile*.

### Inactivation of *camA* reduces sporulation levels *in vitro*

To discover possible functional roles of CamA, we searched among published transposon sequencing studies of *C. difficile*. From a recent study analyzing *C. difficile* gene essentiality and sporulation, we found a homolog of *camA (R20291_2646)* among ~800 genes whose mutation impacted sporulation in the 027 isolate R20291^30^. Given the critical role of sporulation in the persistence and dissemination of *C. difficile* in humans and hospital settings^31^, we decided to test if *camA* inactivation could reduce spore purification efficiencies in the 630Δ*erm* strain background as seen for *R20291_2646* transposon mutants. We used allele-coupled exchange^32^ to construct an in-frame deletion in 630Δ*erm camA* and to complement Δ*camA* with either wild-type *camA* or a variant encoding a catalytic site mutation (N165A) of the MTase in single copy from the *pyrE* locus (Supplementary Fig. 5a; Supplementary Tables 5a, b; Supplementary Materials and Methods; Supplementary Text). We observed that spore purification efficiencies decreased by ~50% in the mutant relative to wild type (Fig. 2a, *P* < 10^−2^, ANOVA, Tukey’s test). Complementation of Δ*camA* with the wild type, but not the catalytic mutant construct, restored spore purification efficiencies to values similar to those observed in wild-type cells (Fig. 2a, Supplementary Table 5c). No differences in growth were observed between wild-type and mutant strains (Supplementary Fig. 5b). Hence, this complementation experiment supports that the loss of methylation events by CamA, rather than the loss of non-catalytic roles of this protein, leads to the decrease in spore yield.

**Fig. 2.**
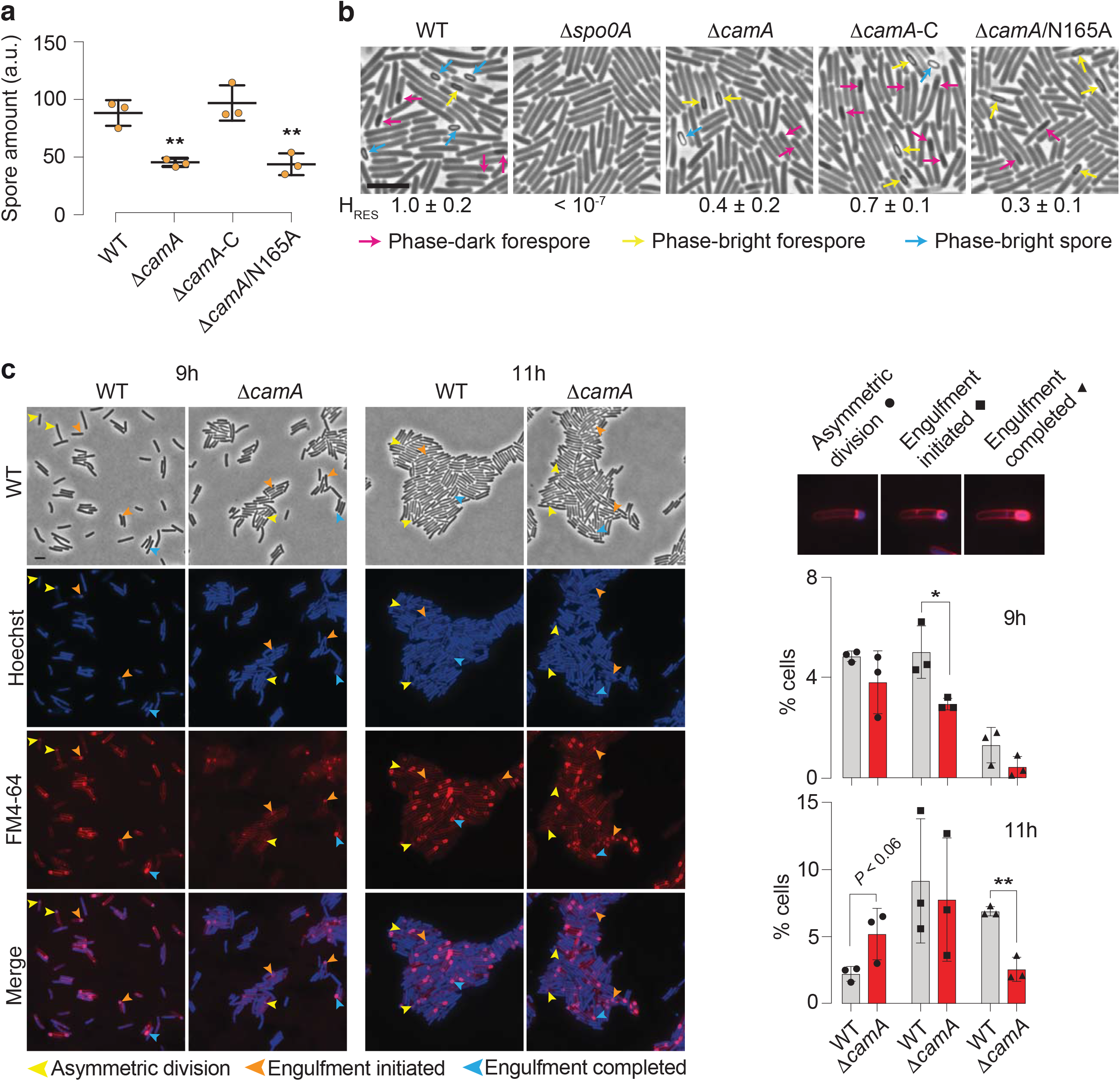
CamA modulates sporulation levels in *C. difficile*. (a) Spore purification efficiencies obtained from sporulating cells harvested 3 days after plating. The spore yield (arbitrary units, a.u.), was determined by measuring the optical density at 600 nm of the resulting spore preparations and correcting for the volume of water in which spores were re-suspended. (b) Phase-contrast microscopy after 20 h of sporulation induction. The Δ*spo0A* strain was used as a negative control because it does not initiate sporulation^2^. Immature phase-dark forespores are marked in pink, and mature phase-bright forespores and free spores are shown in yellow and blue, respectively. Scale bar represents 5 μm. (c) Morphological analysis of wild-type and Δ*camA* cells using fluorescent stains comparing 9 and 11 h following sporulation induction. The polar septum formed during asymmetric division is visible using the FM4-64 membrane stain, while the chromosome that is pumped into the forespore after polar septum formation can be seen as a bright foci using the Hoechst 33342 nucleoid stain^34,88^. FM4-64 staining also allows the engulfing membranes to be visualized. As the mother cell-derived membrane fully encircles the forespore-derived membrane, the FM4-64 signal becomes more intense around the forespore. When these membranes undergo fission, the forespore becomes fully suspended in the mother cell cytosol, and the FM4-64 and Hoechst stains are excluded. Yellow arrows show cells that are undergoing asymmetric division (indicated by a flat polar septum); orange arrows show cells that are in the process of engulfment (indicated by a curved polar septum); and blue arrows show cells that have completed engulfment (indicated by bright membrane staining fully surrounding the forespore). Barplots indicate the number of sporulating cells at different stages of spore assembly in both wild-type and Δ*camA* cells. A minimum of 1000 cells per replicate, per strain, and a minimum of two fields were scored for each strain at each timepoint. * *P* ≤ 0.05, ** *P* < 10^−2^, unpaired Student’s t-test.

The diminished spore purification efficiencies observed in the Δ*camA* mutants could be due to reduced number of cells that induce sporulation or defects in spore assembly^33^. Visual inspection of samples before and after spore purification on a density gradient revealed qualitatively lower levels of mature, phase-bright spores (Supplementary Fig. 5c). Since purified wild-type and Δ*camA* spores had similar levels of chloroform resistance (Supplementary Fig. 5d), and wild-type and Δ*camA* spores germinated with similar efficiency when plated on media containing germinant (Supplementary Fig. 5e), the reduced spore purification efficiencies of the MTase mutants likely reflect a defect in sporulation initiation rather than the sporulation process itself.

Consistent with this hypothesis, fewer Δ*camA* cells were qualitatively observed to be sporulating in phase-contrast microscopy analyses relative to wild type (Fig. 2b). To determine whether loss of CamA impairs the frequency rather than the progression of sporulation, we visualized and quantified the number of sporulating cells at different stages of spore assembly using the FM4-64 membrane stain and Hoechst nucleoid stain^34^ (Fig. 2c).

While similar numbers of wild-type and Δ*camA* cells were observed at asymmetric division (the first morphological stage of sporulation), at the 9 h time point, 50% fewer Δ*camA* cells had initiated engulfment. Furthermore, 11 h after sporulation induction, ~2-fold more Δ*camA* cells were at the asymmetric division stage relative to wild type, whereas 50% fewer Δ*camA* cells had completed engulfment. Since similar numbers of sporulating cells were observed between wild type and Δ*camA* at the 11 h timepoint, Δ*camA’s* sporulation defect appears to arise from fewer cells progressing beyond asymmetric division rather than a defect in sporulation induction.

To confirm that loss of CamA leads to a decrease in the number of cells producing functional spores, we compared the ability of Δ*camA* to form heat-resistant spores capable of germinating and outgrowing using a heat resistance assay^35^. The Δ*camA* mutant and the catalytic mutant complementation strain produced ~50% fewer heat-resistant spores than wild-type and the wild-type complementation strain (Supplementary Fig. 5e, *P* < 10^−3^, ANOVA, Tukey’s test). Taken together, these findings suggest that CAAAA<u>A</u> methylation enhances sporulation *in vitro*. This functional difference prompted us to perform a comprehensive methylome and transcriptome analysis of the wild-type and MTase mutant strains.

### Comparative analysis of CAAAA<u>A</u> sites across *C. difficile* genomes

The *C. difficile* genome has an average of 7,721 (±197, sd) CAAAA<u>A</u> motif sites (Supplementary Table 6a). Adjusted by the k-mer frequency of the AT-rich *C. difficile* genome (70.9%) using Markov models^36^ (Materials and Methods), CAAAA<u>A</u> motif sites are significantly under-represented in the chromosome, particularly in intragenic regions (Supplementary Fig. 6a, Supplementary Table 6b). To evaluate if specific chromosomal regions are enriched or depleted for this motif, we used a multi-scale signal representation (MSR) approach^37^ (Materials and Methods). Briefly, MSR uses wavelet transformation to examine the chromosome at a succession of increasing length scales by testing for enrichment or depletion of a given genomic signal. While scale values <10 are typically associated with regions <100 bp, genomic regions enriched for CAAAA<u>A</u> sites at scale values >20 corresponding to segments larger than 1 kb (*i.e*. gene and operon scale) were observed and included genes related to sporulation and colonization (Fig. 3a, regions A-E; Supplementary Tables 6c, d): stage 0 (*spo0A*), stage III (*spoIIIAA-AH*), and stage IV (*spoIVB, sigK*) of sporulation, membrane transport (PTS and ABC-type transport systems), transcriptional regulation (*e.g*., *iscR, fur*), and multiple cell wall proteins (CWPs).

**Fig. 3.**
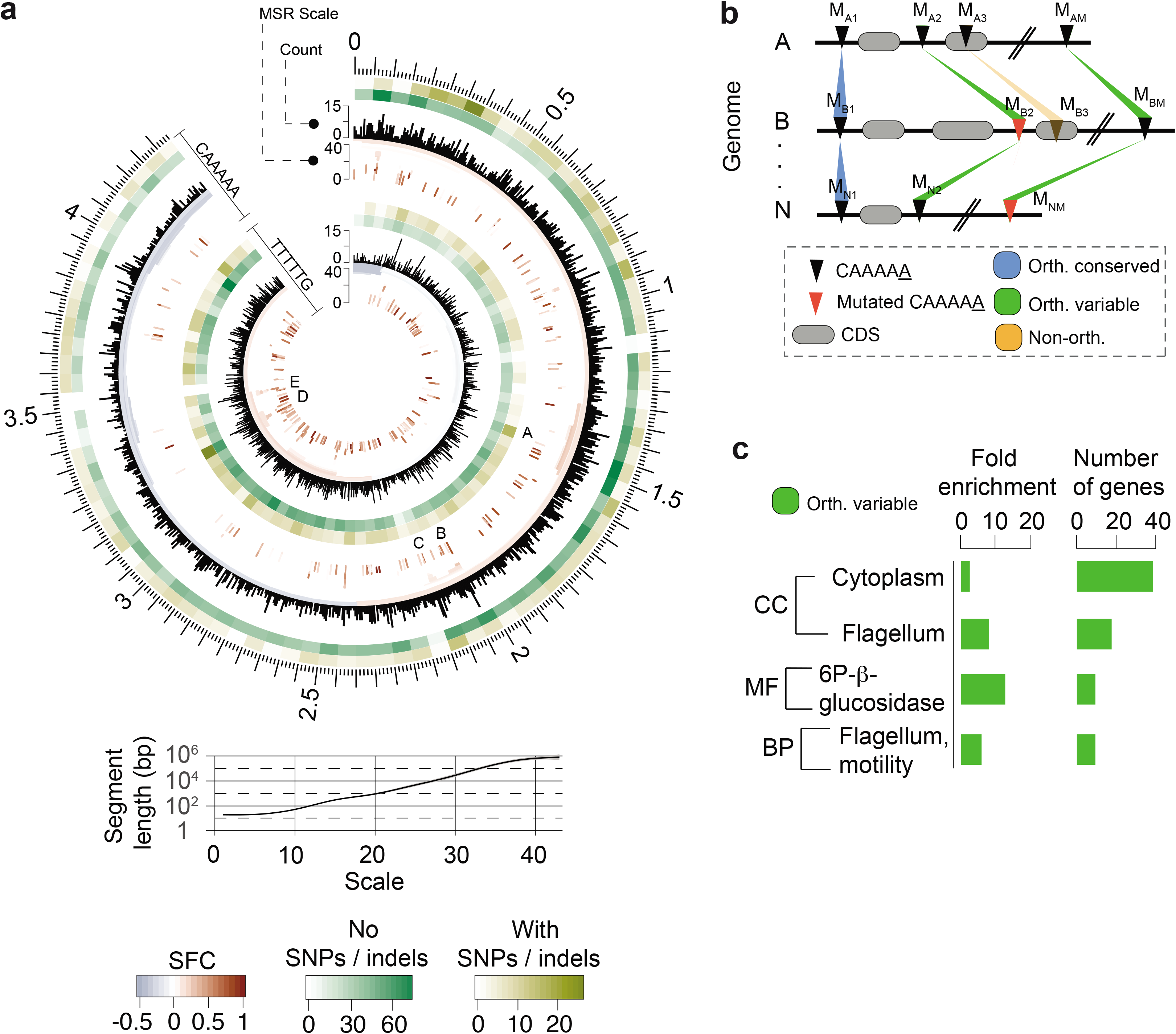
Abundance, distribution, and conservation of CAAAA<u>A</u> motif sites. (a) Distribution of CAAAA<u>A</u> motif sites in both strands of the reference *C. difficile* 630 genome and corresponding genomic signal obtained by MSR. Letters (A-E) represent regions with particularly high abundance of CAAAA<u>A</u> motifs at scales above 20, i.e., typically above the single gene level (median gene size in *C. difficile* = 0.78 kb) (see Supplementary Table 6d for genes contained in each of the regions). Relation between MSR scale and segment length is also shown. The significant fold-change (SFC) corresponds to the fold-change (log_2_ ratio) between observed and randomly expected overlap statistically significant at *P* = 10^−6^. Heatmap layers correspond to the number of orthologous conserved (no SNPs/indels, green-shaded) and orthologous variable (with SNPs/indels) CA_5_ motif positions. (b) Whole genome alignment of 37 *C. difficile* genomes (36 isolates + *C. difficile* 630 as reference) was performed using Mauve. We defined an orthologous occurrence of the CAAAA<u>A</u> motif (black triangles) if an exact match to the motif was present in each of the 37 genomes (conserved, blue-shaded regions), or if at least one motif (and a maximum of *n*-1, being *n* the number of genomes) contained positional polymorphisms (maximum of two SNPs or indels per motif) (variable, green-shaded regions). Non-orthologous occurrences of CAAAA<u>A</u> are indicated as orange-shaded regions. The results are shown in Fig. 3a in the form of heatmaps. Numbering in scheme is based on mapping location. (c) DAVID enrichment analysis of the genes containing intragenic and regulatory (100 bp upstream the start codon) orthologous variable CAAAA<u>A</u> motif sites. We considered a Fisher’s exact test enrichment statistics, a Benjamini-Hochberg corrected *P*-value cutoff of 0.05, and a false discovery rate (FDR) < 0.05.

To further characterize CAAAA<u>A</u> motif sites, we built upon our large collection of *C. difficile* methylomes by categorizing these motif sites on the basis of their positional conservation across genomes. We performed whole genome alignment of 37 *C. difficile* genomes (36 isolates + *C. difficile* 630 as reference) (Materials and Methods, Supplementary Dataset 1), and classified each motif position in the alignment as either: (1) conserved orthologous (devoid of SNPs or indels); (2) variable orthologous (in which at least one genome contains a SNP or indel); and (3) non-orthologous (Fig. 3b). We found a total of 5,828 conserved orthologous motif positions, 1,050 variable orthologous positions (885 with SNPs and 165 with indels), and an average of 843 non-orthologous positions per genome (Supplementary Table 6e). The latter were, as expected, largely mapped to MGEs. Among orthologous positions, the variable ones represent a particularly interesting subset to study, since they contribute to variations of CAAAA<u>A</u> sites across genomes with subsequent methylation abrogation (Supplementary Table 6f). We used DAVID gene enrichment analysis and found cytoplasm- (e.g.: *pheA, fdhD, ogt1*, *spoIVA*) and motility-related genes (e.g.: *fliZ, fliN, fliM, flgL*) to over-represent orthologous variable CAAAA<u>A</u> positions (FDR < 5%, Fig. 3c). Concomitantly, the regions with the highest density of orthologous variable positions were found at the S-layer locus region (region D, Fig. 3a) and between positions 0.31-0.33 Mb, which is rich in flagellar genes. The very large number and dispersion of conserved orthologous positions precluded a similar functional analysis. To test whether homologous recombination (HR) contributes to the cross-genome variation of CAAAA<u>A</u> motif sites located in the core-genome, we also performed a systematic analysis of such events (Supplementary Figs. 6b-d, Supplementary Table 6g, Supplementary Text) and found that HR tracts indeed over-represent (O/E=1.40, *P* < 10^−3^; Chi-square test) orthologous variable CAAAA<u>A</u> motif positions, while the core-genome without HR tracts underrepresents them (O/E=0.89, *P* < 10^−3^; Chi-square test) (Supplementary Figs. 6e, f).

Collectively, genome-wide distribution analyses and across-genome comparative analyses suggest that CAAAA<u>A</u> sites are enriched in regions harboring genes related to sporulation and colonization, orthologous variable CAAAA<u>A</u> positions are enriched in regions harboring genes related to cytoplasm- and motility-related genes, and the former’s variability is at least partially fueled by HR.

### Non-methylated CAAAA<u>A</u> motif sites are enriched in regulatory elements

DNA methylation is highly motif driven in bacteria; i.e. in most cases, >95% of the occurrence of a methylation motif is methylated. However, a small fraction of methylation motif sites can be non-methylated. The on/off switch of DNA methylation in a bacterial cell can contribute to epigenetic regulation as a result of competitive binding between DNA MTases and other DNA binding proteins (e.g. transcription factors, TFs) as previously described for *E. coli*^21,38–40^. Previous bacterial methylome studies analyzing one or few genomes usually have insufficient statistical power to perform a systematic interrogation of non-methylated motifs sites. Building on our rich collection of 36 *C. difficile* methylomes, we performed a systematic detection and analysis of non-methylated CAAAA<u>A</u> sites. To classify a CAAAA<u>A</u> site as non-methylated, we adopted stringent filtering criteria at interpulse duration ratio (ipdR) and sequencing coverage, and found an average of 21.5 non-methylated CAAAA<u>A</u> motif sites per genome (Supplementary Fig. 7a; Supplementary Table 7a). Non-methylated motif sites were found dispersed throughout the full length of the *C. difficile* genome, yet were overrepresented in orthologous variable and non-orthologous CAAAA<u>A</u> positions (O/E=respectively 1.51 and 1.49) and underrepresented in orthologous conserved CAAAA<u>A</u> positions (O/E 0.84) (*P* < 10^−4^; Chi-square test). This is consistent with the idea that variable positions are more likely to be non-methylated to provide breadth of expression variation. Most of the non-methylated positions (85.4% of 245) failed to conserve such status in more than three genomes at orthologous positions (Fig. 4a, Supplementary Table 7a), while a minor percentage of positions (5.5%) remained non-methylated in at least one third of the isolates, suggesting that competitive protein binding is expected to be more active in certain genomic regions (Fig. 4a), *e.g*. upstream of the pathogenicity locus (PaLoc) (position 786,216), in prophage genes (1,593,616), within the *atp* operon (3,430,190), and in the xylose operon (3,561,672) (Supplementary Fig. 7b).

**Fig. 4.**
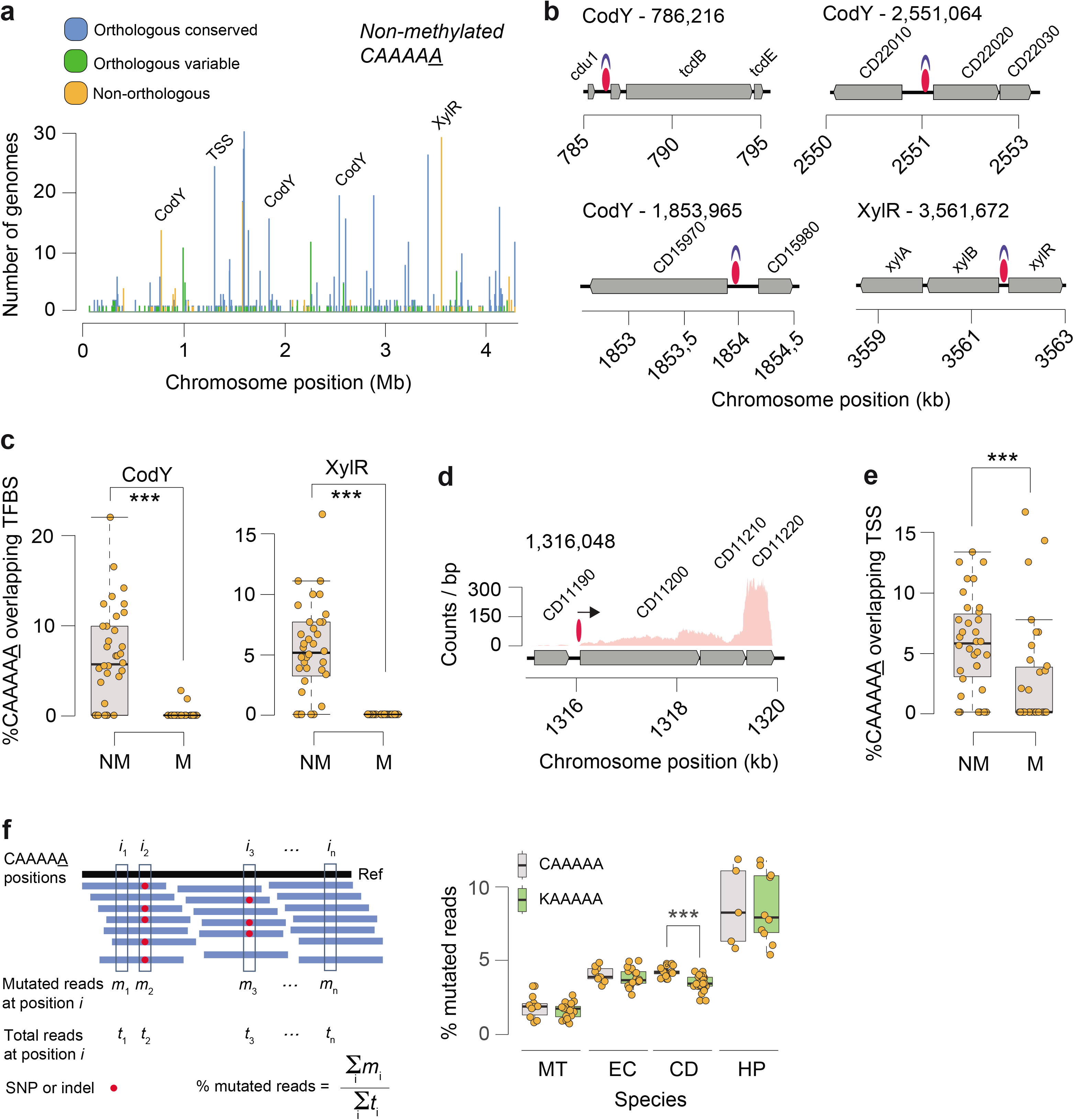
Distribution of non-methylated CAAAA<u>A</u> motif sites, and overlap with transcription factor binding sites (TFBS) and transcription start sites (TSS). (a) Number of *C. difficile* isolates for which non-methylated CAAAA<u>A</u> motif sites were detected at a given chromosome position (coordinates are relative to the reference genome of *C. difficile* 630). Peak colors correspond to orthologous (conserved and variant) and non-orthologous CAAAA<u>A</u> positions. Some of the major peaks of non-methylated CAAAA<u>A</u> positions were found to overlap with TFBS (*e.g*., CodY, XylR) and TSS. (b) Genetic regions for which overlap was observed between highly conserved non-methylated CAAAA<u>A</u> motif sites (red ovals) and TFs (CodY and XylR, shown in blue). Other examples of conserved non-methylated CAAAA<u>A</u> motif sites are illustrated in Supplementary Fig. 7b. (c) % CAAAA<u>A</u> motif sites (non-methylated and methylated) overlapping CodY and XylR for each *C. difficile* isolate. (d) Example of a chromosomal region in which non-methylated CAAAA<u>A</u> motifs overlap a TSS (shown as arrow). (e) % CAAAA<u>A</u> motifs (non-methylated (NM) and methylated (M)) overlapping TSSs for each *C. difficile* isolate. (f) % mutated reads (SNPs + indels) in CAAAA<u>A</u> and K(G or T)AAAAA motifs for *M. tuberculosis* (MT), *E. coli* (EC), *C. difficile* (CD) and *H. pylori* (HP). AAAAAA was not considered as control motif as it would theoretically be more error-prone. *** *P* < 10^−3^, Mann-Whitney-Wilcoxon test.

The non-methylated CAAAA<u>A</u> positions detected across the 36 *C. difficile* genomes allowed a systematic search for evidence of overlap with TF binding sites (TFBSs) and transcription start sites (TSSs). To test this, we queried our genomes for putative binding sites of 21 TFs pertaining to 14 distinct families (Supplementary Table 7b; Materials and Methods) and found overlaps between prominent peaks of non-methylated CAAAA<u>A</u> positions (Fig. 4a) and the TFBSs of CodY and XylR (Fig. 4b, Supplementary Fig. 7b, Supplementary Table 7c). Performing the analysis at the genome level, both CodY and XylR binding sites showed significant enrichment (*P* < 10^−3^, Mann-Whitney-Wilcoxon test) for non-methylated CAAAA<u>A</u> (Fig. 4c; Supplementary Fig. 7c). In a similar enrichment analysis using 2,015 TSSs reconstructed from RNA-seq data coverage^41^ (Materials and Methods; Supplementary Table 7d), we found a genome-level enrichment: non-methylated CAAAA<u>A</u> sites preferentially overlapped with TSSs (Figs. 4d, e; Supplementary Figs. 7d, e; *P* < 10^−3^, Mann-Whitney-Wilcoxon test,).

In addition to the on/off epigenetic switch driven by competitive binding between the MTase and other DNA binding proteins at CAAAA<u>A</u> sites, we hypothesized that epigenetic heterogeneity within a clonal population could also stem from DNA replication errors at CAAAA<u>A</u> sites, especially because homopolymer tracts are expected to be more error-prone^42^. To test this hypothesis, we investigated if CAAAA<u>A</u> had a larger than expected frequency of mutated reads (compared to baseline sequencing errors). Specifically, we tested if CAAAA<u>A</u> sites in *C. difficile* are more error prone during DNA replication, compared to the control motifs TAAAAA or GAAAAA (here called KAAAAA). To account for the confounding effect of mutation rate at different sequence contexts, we further computed the % mutated reads (SNPs + indels) in four distinct species with a broad dispersion of mutation rates (*Mycobacterium tuberculosis* < *E. coli* ≈ *C. difficile* < *Helicobacter pylori)*. Interestingly, we indeed found a significantly higher % of mutated reads mapping to CAAAA<u>A</u> sites compared to KAAAAA sites in *C. difficile* (Fig. 4f, Supplementary Fig. 7f), while this difference was not observed in any of the control genomes. The observation of increased mutagenesis at 6mA target sites is in line with recent data obtained in *Neisseria meningitidis*^43^ and is worth further investigation in future work.

Collectively, these results highlight two types of variations that affect CAAAA<u>A</u> methylation status in *C. difficile:* (i) on/off epigenetic switch of CAAAA<u>A</u> sites preferentially overlapping with putative TFBSs and TSSs, and (ii) higher mutation rates at the CAAAA<u>A</u> motif sites that contribute to cell-to-cell heterogeneity.

### Loss of CAAAA<u>A</u> methylation impacts transcription of multiple gene categories including sporulation

To study the functional significance of methylation at CAAAA<u>A</u> sites, we used RNA-seq to compare the transcriptomes of wild-type *C. difficile* 630Δ*erm* with that of Δ*camA* both in liquid medium (exponential and stationary growth stage) and following sporulation induction (9 and 10.5 h) (Supplementary Fig. 8, Supplementary Table 8a, Materials and Methods). Of the 3,896 genes annotated in *C. difficile* 630, 36 – 361 (0.9 – 9.3%, depending on the time point) were differentially expressed (DE) at a 5% FDR and | log_2_FC | > 1 (2-fold change in gene expression) (Fig. 5a, Supplementary Tables 8b-d). DE genes in Δ*camA* relative to wild type appeared to have a significant enrichment in CAAAA<u>A</u> motif sites compared to non-DE genes (*P* < 10^−2^, Mann-Whitney-Wilcoxon test) in broth culture; a qualitatively similar trend was also observed during sporulation (Supplementary Fig. 9a). Consistent with our finding that loss of CamA reduces spore formation, the transcriptome analyses revealed that 118 and 120 genes previously identified as being induced during sporulation^44,45^ were expressed at ≥ 50% lower levels in Δ*camA* cells relative to wild type at the 9 and 10.5 h timepoints, respectively (Supplementary Table 8b).

**Fig. 5.**
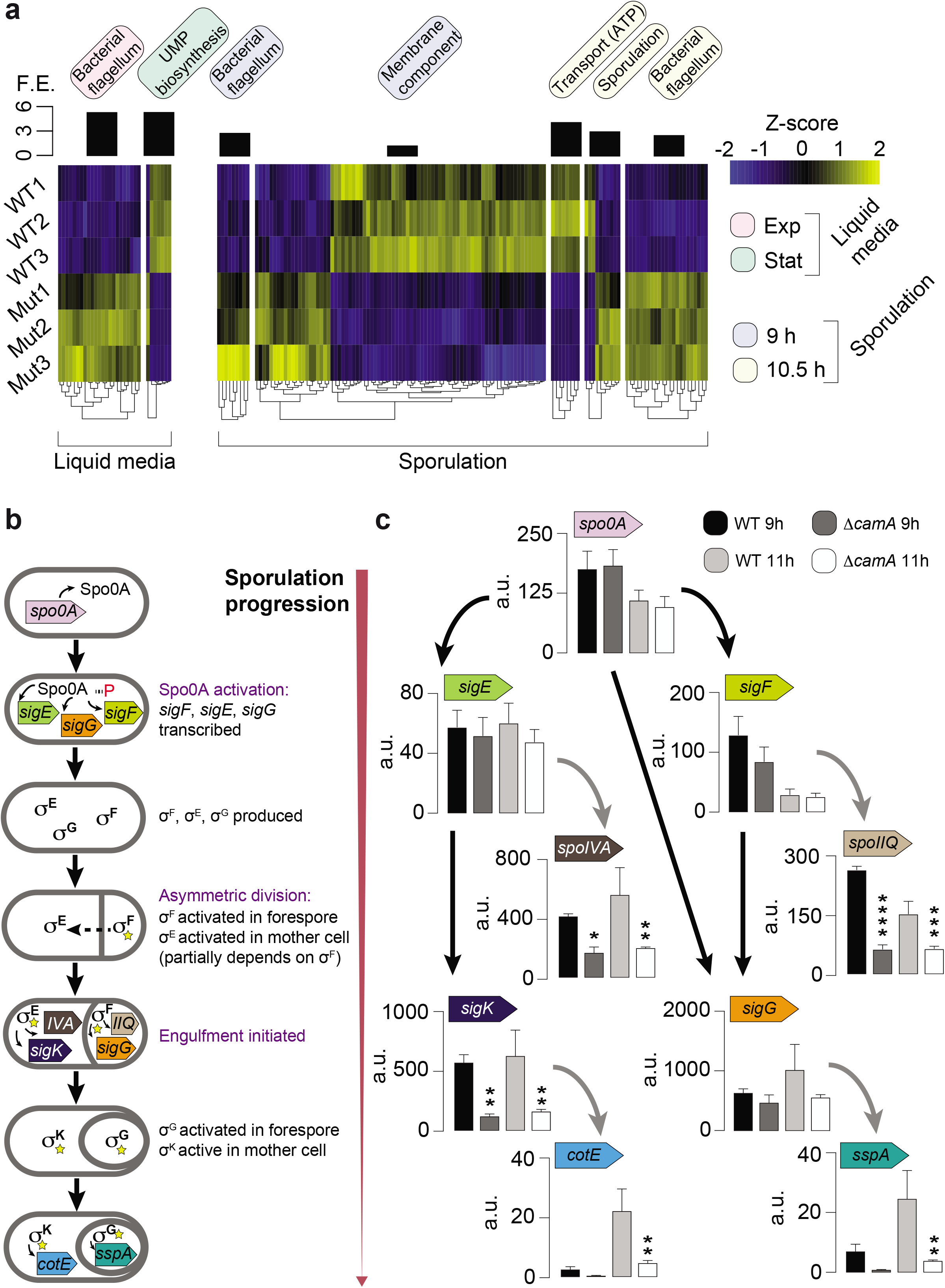
Gene expression analysis. (a) Heatmap of 161 genes in three replicates of *C. difficile* 630 compared to equal number of replicates of *C. difficile* 630Δ*camA* and that are enriched for the GO terms shown in boxes and detailed in Supplementary Table 8c. The *Z* score reflects the degree of down- (*Z* score < 0) or up- (*Z* score > 0) regulation, computed by subtracting the mean of the log-transformed expression values and dividing by the standard deviation for each gene over all samples scored. (b) Schematic illustrating the sequence of sporulation sigma factor gene transcription and protein activation coupled to morphological changes during sporulation. (c) Comparison of relative transcript levels in wild type and Δ*camA* as determined by qRT-PCR for sporulation sigma factor genes and representative genes in the regulons of sporulation-specific sigma factors at 9 and 11 h after sporulation induction. It should be noted that the primers for *sigK* amplify a region prior to the *sigK* excision site^49^. Statistical significance was determined by one-way ANOVA and Tukey’s test for multiple comparisons (* *P* ≤ 0.05, ** *P* < 10^−2^, *** *P* < 10^−3^, **** *P* < 10^−4^).

The transcriptional program that mediates sporulation in *C. difficile* is controlled by a master transcriptional activator, Spo0A, and four sporulation-specific sigma factors, σ^F^, σ^E^, σ^G^ and σ^K^. These factors activate distinct regulons that ultimately lead to the assembly of functional spores^37,38^ (Fig. 5b). Activated Spo0A induces the expression of genes encoding σ^F^, σ^E^, and σ^G^ as well as factors required for asymmetric division and the post-translational activation of the early-stage sporulation sigma factors, σ^F^ and σ^E^ (Fig. 5b). σ^F^ is the first sporulation-specific sigma factor to be fully activated, and it only becomes active in the forespore after asymmetric division is completed^46^. Activated σ^F^ subsequently induces the transcription of genes whose products mediate σ^G^ activation in the forespore and partially mediates σ^E^ activation in the mother cell^47^. Activated σ^E^ induces the transcription of *sigK*^48^ and factors required for the excision of a prophage-like element from the *sigK* gene^49^. Thus, *C. difficile* sporulation is controlled by a transcriptional hierarchy that is coupled to morphological events such that downstream sigma factors (σ^G^ and σ^K^) depend on the activation of upstream sigma factors (σ^F^ and σ^E^). Notably, genes in the regulons of all four sporulation-specific sigma factors were under-expressed in Δ*camA* relative to wild type, whereas a relatively small subset of Spo0A regulon genes exhibited this pattern of regulation (Supplementary Fig. 9b, Supplementary Table 8e). These observations suggested that loss of CamA reduces spore formation by targeting events early on during sporulation.

To gain further insight into the regulatory stage of sporulation that CamA-mediated DNA methylation specifically impacts, we used qRT-PCR to analyze the expression of genes encoding Spo0A, the sporulation-specific sigma factors^44,50^, and their individual regulons^51–53^. These analyses were conducted on a separate set of RNA samples harvested at 9 and 11 h following sporulation induction. Consistent with our RNA-Seq analyses, Spo0A regulon genes, *spo0A, sigF*, and *sigE*^44,51^, were expressed at similar levels between wild type and Δ*camA* at both 9 and 11 h, implying that the Δ*camA* mutant activates Spo0A at levels similar to wild type. In contrast, σ^F^ and σ^E^ regulon genes, *spoIIQ* and *spoIVA*^44,54^, respectively, were under-expressed in Δ*camA* relative to wild type (Fig. 5b). Reduced SpoIIQ and SpoIVA levels were observed in Δ*camA* by western blot, confirming the transcriptional analyses (Supplementary Fig. 9c). Based on the hierarchical organization of the sporulation regulatory cascade, these results implicate σ^F^ activation as the earliest sporulation stage affected by CamA. This conclusion is supported by our morphological analyses, since fewer Δ*camA* cells initiate and complete engulfment relative to wild type, and engulfment requires both σ^F^- and σ^E^-dependent gene products^55^. Notably, similar numbers of Δ*camA* and wild-type cells are sporulating based on these morphological analyses (Fig. 2c), consistent with the similar levels of *spo0A* observed in WT and Δ*camA* (Fig. 5b). While a small subset of Spo0A regulon genes are under-expressed in Δ*camA* cells, at least some of these genes are dually regulated by Spo0A and σ^F^. For example, transcription of *spoIIR*^52^, which encodes a signaling protein required for σ^E^ activation, is activated by both Spo0A and σ^F^ ^44,47^.

### *In vivo* impacts of the *camA* mutation

To test whether the sporulation defect of Δ*camA* impacts *C. difficile* infection or transmission, we analyzed the effect of the Δ*camA* mutation in an established mouse model of infection in which a cocktail of antibiotics is used to sensitize the animals to *C. difficile* infection^56^. When inoculated with 630Δ*erm* strains, antibiotic-treated mice typically do not develop fulminant disease and instead serve as a model of intestinal colonization and persistence by *C. difficile*^57,58^. Groups of mice (6 males, 6 females) were inoculated by oral gavage with 10^5^ spores of the three genotypes: wild type, Δ*camA*, and Δ*camA-C*. No mortality was observed at the given doses of *C. difficile* spores as expected. Fecal samples were collected every 24 h for seven days. All three *C. difficile* strains reached comparable levels in feces at days 1 and 2 post-inoculation, indicating that they germinate and establish colonization equally (Fig. 6a, Supplementary Materials and Methods). As expected, CFU levels decreased steadily from day 2 post-inoculation to day 7. However, the Δ*camA* mutant showed CFU levels 10-100 times lower than those observed in the wild-type and complemented strains throughout this time frame (ANOVA, *P* < 10^−4^). The bacteria declined below the limit of detection in the feces 6 days post-inoculation for the MTase mutant, while they remained detectable at days 6 and 7 for the wild-type and complemented strains.

**Fig. 6.**
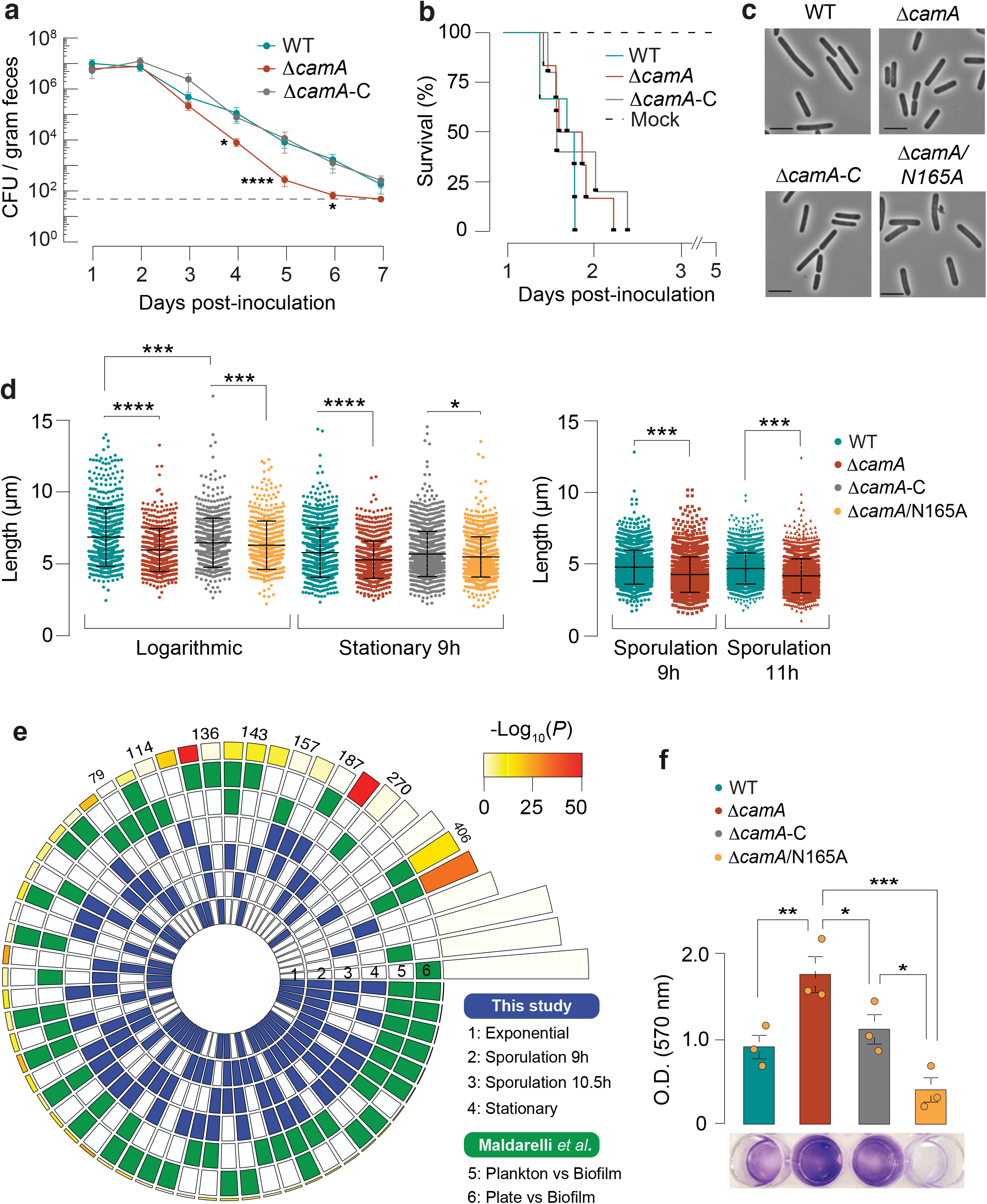
*In vivo* and additional functional impacts of the Δ*camA* mutation. (a) Kinetics of infection in mice (*n*=12) following inoculation with a sub-lethal amount (10^5^ spores) of wild type (WT) *C. difficile* 630Δ*erm*, MTase mutant Δ*camA*, and complement Δ*camA-C*. Dotted line indicates the limit of detection. Log_10_-transformed data from each time point were analyzed by ANOVA. * *P* < 0.05, **** *P* < 10^−4^. (b) Kaplan-Meier survival curves for clindamycin-treated golden Syrian hamsters (n=6) infected with 10^3^ spores of either wild type (WT) *C. difficile 630Δerm*, MTase mutant Δ*camA*, and complement Δ*camA-C*. (c) Representative phase-contrast images of vegetative wild type, Δ*camA*, Δ*camA-C*, and Δ*camA/N165A*, grown in BHIS broth (scale bar 5 μm). (d) Comparison of cell length. A minimum of 400 cells were measured for each strain at mid-log and stationary phase, and a minimum of 1,600 cells were analyzed during sporulation at the indicated timepoints. All cell length measurements were conducted on three independent biological replicates. Statistical significance was determined by one-way ANOVA and Tukey’s test for multiple comparisons. (e) Significance of overlap between multiple datasets of DE genes. Comparisons were performed between DE genes called in this study for each time point (blue-shaded) and those from Maldarelli *et al.*^59^ (green-shaded). The latter corresponds to *C. difficile* DE genes in conditions favoring biofilm formation compared to growth on a plate or planktonic form. Color intensities of the outermost layer represent the *P*-value significance of the intersections (3,896 genes were used as background). The height of the corresponding bars is proportional to the number of common genes in the intersection (indicated at the top of the bars for pairwise comparisons between the different studies). (f) Biofilm production as measured by crystal violet staining absorbance at 570 nm. Error bars correspond to standard error of the mean of three independent experiments, with each strain assayed in quadruplicate in each experiment. * *P* < 0.05, ** *P* < 10^−2^, *** *P* < 10^−3^, unpaired Student’s t-test.

To test whether loss of CamA leads to defects in virulence, we compared Δ*camA* and wild type in a hamster model of infection. Clindamycin-treated golden Syrian hamsters are highly susceptible to the effects of the *C. difficile* toxins and thus represent a model of acute disease. Groups of 6 hamsters (3 male, 3 female) were inoculated by oral gavage with 10^3^ spores of the wild-type, Δ*camA*, and Δ*camA-C* strains. These *C. difficile* strains elicited diarrheal symptoms and weight loss in this model, and we observed no difference in animal survival times post inoculation (Fig. 6b, Supplementary Materials and Methods). This result is consistent with the observation that the wild-type, Δ*camA*, and Δ*camA-C* strains exhibit no differences in toxin gene expression (Supplementary Table 8a) and produce comparable levels of TcdA *in vitro* (Supplementary Fig. 9d). Together, these data indicate that CAAAA<u>A</u> methylation by CamA does not influence toxin-mediated aspects of *C. difficile* pathogenesis but instead impacts *C. difficile’s* ability to persist within the host intestinal tract.

### Additional functional impacts of the *camA* mutation

Considering the high conservation of *camA* across *C. difficile* genomes, we asked if some additional phenotypes could be impacted by the gene’s inactivation. While analyzing images of sporulating *C. difficile*, we noticed that Δ*camA* mutant cells appeared to be shorter on average than wild-type cells. To test this possibility, we measured the lengths of wild-type and Δ*camA* cells during log-phase and stationary-phase growth in broth culture and during sporulation (9 and 11 h time points). These analyses revealed that Δ*camA* cells were ~15% shorter than wild-type cells (*P* < 0.001, one-way ANOVA, Fig. 6c, d) even though no difference in cell growth was observed (Supplementary Fig. 5b). Interestingly, genes encoding putative cell wall remodeling enzymes, *spoVD, cwp16*, and *cwp17*, were over-expressed in the Δ*camA* mutant relative to wild type during growth in broth culture (Supplementary Fig. 9e).

In addition to the above experimental characterization, we attempted to build on the RNA-seq data further beyond the sporulation phenotype, with a special focus on biological processes critical to *C. difficile* infection. Specifically, we performed an overlap analysis between the list of DE genes from our RNA-seq data (wild type vs. Δ*camA* mutant; four different time points) and those from published studies focusing on the colonization and infection by this pathogen (Materials and Methods). Out of the total 20 pairwise overlap analyses, we observed 9 significant overlaps (*P* < 0.01 with Bonferroni correction) between our dataset and those of others (Supplementary Table 8f). First, DE genes in the Δ*camA* mutant (sporulation phases) had a significant overlap to DE genes in conditions favoring the production of biofilm on a solid substrate^59^ (Fig. 6e, Supplementary Table 8f) (O/E=1.3-1.6, *P* < 10^−9^, Chi-Square test). Motivated by this significant overlap, we performed biofilm assays using crystal violet staining of adherent biofilm biomass, and made consistent observation that the Δ*camA* mutant produced more biofilm than wild type (Fig. 6f). These results suggest that methylation inhibits the expression of genes that promote biofilm formation, *e.g*., encoding factors involved in the adherence to a surface and/or stabilizing interbacterial interactions. The differences in biofilm production between Δ*camA* and Δ*camA/N165A* could be explained if the latter retained some DNA binding ability capable of altering transcription of some genes even in the absence of methylation. Second, by comparing DE genes in Δ*camA* relative to wild type with genes DE during infection in different murine gut microbiome compositions^60^, we found a statistically significant overlap, particularly genes expressed at stationary stage (O/E=1.4, *P* < 10^−6^, Chi-Square test) (Supplementary Fig. 10a, Supplementary Table 8f). Lastly, using DE genes obtained from murine gut isolates at increasing time points after infection^61^, we found significant overlaps with DE genes in the Δ*camA* mutant (O/E=1.5, *P* < 10^−4^, Chi-Square test) (Supplementary Fig. 10b, Supplementary Table 8f). Collectively, these integrative overlap analyses provide additional evidence that DNA methylation events by CamA may directly and/or indirectly affect the expression of multiple genes involved in the *in vivo* colonization and biofilm formation of *C. difficile* and inspire future work to elucidate the mechanisms underlying the functional roles of CAAAA<u>A</u> methylation in *C. difficile* pathogenicity.

## Discussion

*C. difficile* is responsible for one of the most common hospital-acquired infections and classified by the US Centers for Disease Control and Prevention as an urgent healthcare risk with significant morbidity and mortality^9^. Because CDI is spread by bacterial spores found within feces, extensive research has been devoted to better understand the genome of this critical pathogen and its sporulation machinery. To address these common goals, we performed the first comprehensive characterization of the DNA methylation landscape across a diverse collection of clinical isolates. During our epigenome analysis, we identified an 6mA MTase (*camA*) conserved across all isolates (and in another ~300 published *C. difficile* genomes) sharing a common methylation motif (CAAAA<u>A</u>). Inactivation of the gene encoding this MTase resulted in a sporulation defect *in vitro* (Fig. 2). This represents, to the best of our knowledge, the first time that DNA methylation has been found to impact sporulation in any bacterium, opening a new dimension of study in *C. difficile*. Infection studies using the mouse model indicate a role for CamA in the persistence of *C. difficile* in the intestinal tract. Since enumeration of *C. difficile* recovered in feces of the infected animals reflects the number of *C. difficile* spores in the gut, the reduced burden of Δ*camA* in the mouse may be due to the mutant’s defect in sporulation, as the ability to form spores was previously shown to be important for persistence^2^. That Δ*camA* exhibited virulence comparable to the wild type in the hamster model suggests that DNA methylation does not impact toxin-mediated disease. However, due to the pleiotropic nature of the MTase it remains possible that multiple factors contribute to the more pronounced effect observed in the mouse model.

The highly conserved *camA* and its flanking genes across *C. difficile* genomes suggest that additional phenotypes may be regulated by CamA beyond sporulation. Consistently, CAAAA<u>A</u> sites were overrepresented in a set of regions enriched in genes with functions linked to sporulation, motility, and membrane transport. Further supporting a broader regulatory network of CamA, is that its loss reduces cell length and results in statistically significant overlap between transcriptional signatures identified in our study (wild type vs Δ*camA* mutant) and those of others observed during the *in vivo* colonization and biofilm formation (Fig. 5b, Supplementary Figs. 7b, c).

The fact that *camA* is a solitary MTase gene without a cognate restriction gene further supports a view that widespread methylation in bacteria has a functional importance beyond that attributed to R-M systems. Previously, the most extensively characterized 6mA MTase was Dam targeting G<u>A</u>TC in *E. coli*. Dam plays multiple important functions and is essential in some pathogens^21^. However, since it is conserved in the large diversity of γ-proteobacteria, it was not considered a promising drug target. In contrast, the uniqueness of *camA* in all *C. difficile* genomes and in just a few *Clostridiales* makes it a promising drug target that may inhibit *C. difficile* in a much more specific manner, which is particularly relevant since gut dysbiosis potentiates *C. difficile* infection^62,63^. In addition, since this MTase seemingly does not impact the general fitness of *C. difficile*^30^, a drug specifically targeting it may be developed with a lower chance for resistance.

Considering the large number of genes differentially expressed in the Δ*camA* mutant, the functional impact of CAAAA<u>A</u> methylation is likely mediated by multiple genes that are either directly regulated by DNA methylation or indirectly regulated by a transcriptional cascade. Mechanistically, DNA methylation can either activate or repress a gene depending on other DNA binding proteins that compete with DNA MTases^16,17,21,64^, so the competition between transcription factors and MTases may form an epigenetic switch to turn on/off a gene.

With more than 2,200 bacterial methylomes published to date, it is becoming increasingly evident that epigenetic regulation of gene expression is highly prevalent across bacterial species. Despite the exciting prospects for studying epigenetic regulation, our ability to comprehensively analyze bacterial epigenomes is limited by a bottleneck in integratively characterizing methylation events, methylation motifs, transcriptomic data, and functional genomics data. In this regard, this work represents the first comprehensive comparative analysis of a large collection of a single bacterial species and thus provides a detailed roadmap that can be used by the scientific community to leverage the current status quo of epigenetic analyses.

## Materials and Methods

### Data and code availability

Genome assemblies are available via NCBI under BioProject ID PRJNA448390. RNA-Seq data has been submitted to the NCBI Sequence Read Archive (SRA) under project PRJNA445308. Scripts and a tutorial supporting all key analyses of this work will be made publicly available at http://github.com/fanglab/ upon the acceptance of the manuscript.

### *Clostridium difficile* isolates and culture

36 clonal C. difficile isolates from CDI fecal samples were obtained using protocols developed in an ongoing Pathogen Surveillance Program at Mount Sinai Hospital (Supplementary Table 1). Additionally, 9 fully sequenced and assembled *C. difficile* genomes were retrieved from Genbank Refseq (ftp://ftp.ncbi.nih.gov/genomes, last accessed in November 2016) (Supplementary Table 1). Raw sequencing data from global and UK collections comprising 291 *C. difficile* 027/BI/NAPI genomes were used^11^ (Supplementary Table 4). *C. difficile* positive stool samples were frozen at −80 °C prior to analysis. All stool samples underwent culture for *C. difficile* using ethanol shock culture method, adapted from Griffiths et al. Briefly, approximately 80 mg of solid stool (50 μl liquid stool samples) was added to 0.5 ml of 70% ethanol wash and the sample was vortex mixed and incubated at room temperature for 20 min. A loopful was then cultured onto *C. difficile* selective agar (CDSA, Becton Dickinson, Franklin Lakes, NJ) and the plates were incubated anaerobically at 37 °C for up to 72 h. A single colony was subcultured onto a Trypticase™ soy agar with 5% defibrinated sheep blood plate (TSA II™, Becton Dickinson, Franklin Lakes, NJ) and incubated anaerobically at 37 °C for 48 h, after which colonies giving the characteristic *C. difficile* odor and fluorescence under UV illumination were obtained and confirmed by MALDI on a Brucker biotyper. For long-term storage, individual colonies were emulsified in tryptic soy broth containing 15% glycerol and stored at −80 °C.

### Single-molecule real-time (SMRT) sequencing

Primer was annealed to size-selected (>8 kb) SMRTbells with the full-length libraries (80 °C for 2 min and 30 s followed by decreasing the temperature by 0.1 °C increments to 25 °C). The polymerase-template complex was then bound to the P6 enzyme using a ratio of 10:1 polymerase to SMRTbell at 0.5 nM for 4 h at 30 °C and then held at 4 °C until ready for magbead loading, prior to sequencing. The magnetic bead-loading step was conducted at 4 °C for 60 min per manufacturer’s guidelines. The magbead-loaded, polymerase-bound, SMRTbell libraries were placed onto the RSII machine at a sequencing concentration of 125-175 pM and configured for a 240 min continuous sequencing run.

### *De novo* genome assembly and motif discovery

The RS_HGAP3 protocol was used for *de novo* genome assembly, followed by custom scripts for genome finishing and annotation. RS_Modification_and_Motif_Analysis.1 was used for *de novo* methylation motif discovery. A custom script was used to examine each motif to ensure its reliable methylation states. In brief, variations of a putative motif are examined by comparing the ipdR distribution of each variation with non-methylated motifs.

### Identification of R-M systems

Identification of R-M systems was performed as previously described^14^. Briefly, curated reference protein sequences of Types I, II, IIC and III R-M systems and Type IV REases were downloaded from the data set ‘gold standards’ of REBASE^65^ (last accessed in November 2016). All-against-all searches were performed for REase and MTase standard protein sequences retrieved from REBASE using BLASTP v2.5.0+ (default settings, *e* value < 10^−3^). The resulting *e* values were log-transformed and used for clustering into protein families by Markov Clustering (MCL) v14-137^66^. Each protein family was aligned with MAFFT v7.305b^67^ using the E-INS-i option, 1,000 cycles of iterative refinement, and offset 0. Alignments were visualized in SEAVIEW v4.6.1^68^ and manually trimmed to remove poorly aligned regions at the extremities. Hidden Markov model (HMM) profiles were then built from each multiple sequence alignment using the hmmbuild program from the HMMER v3.0 suite^69^ (default parameters) (available at https://github.com/pedrocas81). Types I, II, and III R-M systems were identified by searching genes encoding the MTase and REase components at less than five genes apart.

### CAAAA<u>A</u> motif abundance and exceptionality

We evaluated the exceptionality of the CAAAA<u>A</u> motif using R’MES^36^ v3.1.0 (http://migale.jouy.inra.fr/?q=rmes). This tool computes scores of exceptionality for k-mers of length *l*, by comparing observed and expected counts under Markov models that take sequence composition under consideration. R’MES outputs scores of exceptionality, which are, by definition, obtained from *P*-values through the standard one-to-one probit transformation. Analysis of motif abundance was performed with a previous developed framework^37^ involving a multi-scale representation (MSR) of genomic signals. We created a binary genomic signal for motif content, which was 1 at motif positions, and 0 otherwise. 50 length scales were used. Pruning parameter values were set to default and the *P*-value threshold to 10^−6^.

### Whole-genome multiple alignment and classification of CAAAA<u>A</u> positions

Whole-genome multiple alignment of 37 genomes (36 *C. difficile* isolates and *C. difficile* 630) was produced by the progressiveMauve program^70^ v2.4.0 with default parameters. Since progressiveMauve does not rely on annotations to guide the alignment, we first used the Mauve Contig Mover^71^ to reorder and reorient draft genome contigs according to the reference genome of *C. difficile* 630. A core alignment was built after filtering and concatenating locally collinear blocks (LCBs) of size ≥50 bp using the stripSubsetLCBs script (http://darlinglab.org/mauve/snapshots/2015/2015-01-09/linux-x64/). The lower value chosen for LCB size accounts for the specific aim of maximizing the number of orthologous motifs detected. The XMFA output format of Mauve was converted to VCF format using dedicated scripts, and VCFtools^72^ was used to parse positional variants (SNPs and indels). Orthologous occurrences of the CAAAA<u>A</u> motif were defined if an exact match to the motif was present in each of the 37 genomes (conserved orthologous positions), or if at least one motif (and a maximum of *n*-1, with *n* being the number of genomes) contained positional polymorphisms (maximum of two SNPs or indels per motif) (variable orthologous positions). Non-orthologous occurrences of CAAAA<u>A</u> were obtained from the whole genome alignment before the extraction of LCBs. The former correspond to those situations where the CAAAA<u>A</u> motif was absent in at least one genome. Typically, these correspond to regions containing MGEs or unaligned repetitive regions.

### Identification of transcription factor binding sites, and transcription start sites

Identification of transcription factor binding sites (TFBS) was performed by retrieving *C. difficile* 630 regulatory sites in FASTA format from the RegPrecise database (http://regprecise.lbl.gov^73^, last accessed July 2017). These were converted to PWMs using in-house developed scripts. This led to a total of 21 PWMs pertaining to 14 distinct transcription factor families (Supplementary Table 7b). Matches between these matrices and *C. difficile* genomes was performed with MAST^74^ (default settings). MAST output was filtered on the basis of *P*-value. Hits with *P*-value <10^−9^ were considered positive, while hits >10^−5^ were considered negative. Hits with intermediate *P*-values were only considered positive if the *P*-value of the hit divided by the *P*-value of the worst positive hit was lower than 100. For the CcpA, LexA, NrdR, and CodY (which have shorter binding sites), we considered positive hits those with *P*-values <10^−8^. Transcription start sites (TSSs) were predicted with Parseq^41^ under the ‘fast’ speed option from multiple RNA-seq datasets (see below). Transcription and breakpoint probabilities were computed using a background expression level threshold of 0.1 and a score penalty of 0.05. We kept only high-confidence 5’ breakpoint hits, located at a maximum distance of 200 bp from the nearest start codon. A ±5 bp window around the TSS was considered if only one single predicted value was obtained; otherwise we considered an interval delimited by the minimum and maximum values predicted by Parseq.

### RNA sequencing, read alignment, and differential expression analysis

Purified RNA was extracted from three biological replicates of sporulating (9, 10.5 h) and exponential and stationary grown cultures of *C. difficile* 630Δ*erm* and *C. difficile* 630Δ*erm*Δ*camA*, DNase-treated, ribosomal RNA-depleted, and converted to cDNA as previously described^44^ (more details in Supplementary Methods). RNA sequencing was performed on a HiSeq 2500, yielding an average of 29.4 (±4.5, sd) million 100-bp single-end reads per sample (exponential and stationary growth timepoints) and 26.9 (±4.3, sd) million 150-bp paired-end reads per sample (sporulation time points). Read quality was checked using FastQC v0.11.5 (http://www.bioinformatics.babraham.ac.uk/projects/fastqc). We used Trimmomatic^75^ v0.39 to remove adapters and low-quality reads (parameters: PE, -phred33, ILLUMINACLIP:<adapters.fa>:2:30:10:8:True, SLIDINGWINDOW:4:15, LEADING:20 TRAILING:20, MINLEN:50). Subsequently, rRNA sequences were filtered from the data set using SortMeRNA^76^ v2.1, based on the SILVA 16s and 23s rRNA databases^77^, and Rfam 5s rRNA database^78^. The resulting non-rRNA reads were mapped to the *C. difficile* 630 reference genome using BWA-MEM v0.7.17-r1198^79^. The resulting bam files were sorted and indexed with Samtools^80^ v1.9, and read assignment was performed with featureCounts^81^ v1.6.4 (excluding multi-mapping and multi-overlapping reads). A gene was included for differential expression analysis if it had more than one count in all samples. Normalization and differential expression testing were performed using the Bioconductor package DESeq2 v1.18.1^82^. Genes with a false discovery rate (FDR) < 0.05 and |log_2_FC| > 1 were called as differentially expressed. Functional classification of genes was performed using the DAVID online database (https://david.ncifcrf.gov)^83^ with Fisher’s exact test enrichment statistics, a Benjamini-Hochberg corrected *P*-value cutoff of 0.05, and FDR < 0.05. The reproducibility of DAVID’s functional classification was tested with Blast2GO v5.2^84^ and Panther v14^85^. Briefly, for Blast2GO, we ran BLASTX searches of the *C. difficile* 630 genome against the entire GenBank bacterial protein database (as of 09/2018). The output, in XML format, was loaded into Blast2GO, and mapping, annotation and enrichment analysis was performed as indicated (http://docs.blast2go.com/user-manual/quick-start/). For Panther, we downloaded the most recent HMM library (ftp.pantherdb.org/hmm_scoring/13.1/PANTHER13.1_hmmscoring.tgz), and annotated our *C. difficile* 630 protein set with pantherScore2.1.pl. Both input and background gene lists were formatted to the Panther Generic Mapping File type, as described in the website (http://www.pantherdb.org). To assess the significance of the intersection between multiple datasets of differentially expressed genes (typically observed during *C. difficile* colonization and infection), we collected gene-expression data from *in vivo* and *in vitro* studies^59–61^, in which key factors for gut colonization (*e.g*., time post-infection, antibiotic exposure, and spatial structure (planktonic, biofilm growth)) were tested. Differentially expressed genes were called under the same conditions as described above. Statistical analyses and graphical representation of multi-set intersections was performed with the R package *SuperExactTest*^86^.

## Acknowledgements

We acknowledge Dr. Richard J. Roberts (New England Biolabs, Inc. USA) for his expertise and generous help with the prediction of R-M systems and orphan MTases in *C. difficile* genomes using REBASE Tools, his critical reading, and for providing helpful comments/suggestions. He was originally an author of this manuscript, however, as a staunch supporter of the open access movement, he will not author a paper that is not open access We also acknowledge Dr. Eduardo P.C. Rocha (Institut Pasteur, Paris, France) for critical reading and for providing helpful comments/suggestions. The work was primarily funded by R01 GM114472 (G.F.) from the National Institutes of Health and Icahn Institute for Genomics and Multiscale Biology. In addition, the work was funded by NIH grants R01 AI119145 (H.v.B and A.B.), R01 AI22232 (A.S.) and R01 AI107029 (R.T.) a Hirschl Research Scholar award from the Irma T. Hirschl/Monique Weill-Caulier Trust (G.F.), a Pew Scholar in the Biomedical Sciences grant from the Pew Charitable (A.S.). G.F. is a Nash Family Research Scholar. A.S. holds an Investigators in the Pathogenesis of Infectious Disease Award from the Burroughs Wellcome Fund. This work was also supported in part through the computational resources and staff expertise provided by the Department of Scientific Computing at the Icahn School of Medicine at Mount Sinai.

## Author Contributions

G.F. conceived the hypothesis. A.S. and G.F. supervised the project. P.H.O. and G.F. designed the computational methods. P.H.O., R.T., A.S., and G.F. designed the experiments. P.H.O. performed most of the computational analyses and developed most of the scripts supporting the analyses. J.W.R. performed the growth curves, microscopy analyses (fluorescence and phase-contrast), analyses of cell length and sporulation stage, isolated some of the RNA and processed it for qRT-PCR studies; qRT-PCR analyses of sporulation genes. A.S. constructed the deletion and catalytic Δ*camA* mutants, performed complementation, isolated and processed the RNA for several of the RNA analyses and performed many of the sporulation phenotypic assays. E.M.G. and D.T. performed the animal infection experiment and analyzed the data under the supervision of R.T. A.Kim and G.F. performed methylation motif discovery and refinement. O.S. and E.A.M. performed qRT-PCR controls for RNA-seq analyses. O.S., E.A.M., G.D., M.L., C.B., N.Z., D.A., I.O., G.P., C.H., S.H., R.S., H.v.B. and A.S. contributed to the other experiments. G.D., I.O., and R.S. designed and conducted SMRT sequencing. P.H.O., J.W.R., E.M.G., D.T., A.Kim., O.S., T.P., S.Z., E.A.M., M.T., S.B., A.A., A.B., R.J.B., R.T., E.E.S., R.S., H.B., A.Kasarskis., R.T., A.S. and G.F. analyzed the data. P.H.O., R.T., A.S., and G.F. wrote the manuscript with additional information inputs from other co-authors.

