## Supplementary Information for "Epigenomic and functional characterization of a core DNA methyltransferase in the human pathogen *Clostridium difficile*"

#### Inventory of Supplementary information

**Supplementary Text**

**Supplementary Materials and Methods**

**Supplementary Fig. 1.** Multiple defense systems and gene flux control in *C. difficile*.

**Supplementary Fig. 2.** Relation between gene flux and CRISPR spacer content.

**Supplementary Fig. 3.** Interplay between Type I R-M systems and gene flux in *C. difficile*.

**Supplementary Fig. 4.** Genomic context and conservation of *camA*.

**Supplementary Fig. 5.** Δ*camA* construction, purified spore analyses, and sporulation kinetics.

**Supplementary Fig. 6.** Core- / Pan-genome analyses of *C. difficile* and HR landscape.

**Supplementary Fig. 7.** Non-methylated CAAAA<u>A</u> motif sites. Motif overlap with TFBS and TSS.

**Supplementary Fig. 8.** Principal Component Analysis (PCA) and MA-plots for RNA-seq data.

**Supplementary Fig. 9.** DE, gene, and protein expression analyses.

**Supplementary Fig. 10.** Overlap between multiple datasets of DE genes.

### Supplementary Text

#### 5mC methylation motifs

While SMRT-seq can effectively detect 6mA and 4mC events, it does not effectively detect 5mC events[1]. Therefore, no confident 5mC motifs were detected from SMRT-seq data. Because MTase gene analysis suggests that <10% of MTases are of the 5mC type across the 36 *C. difficile* genomes, we focused on 6mA and 4mC methylation in this study.

#### A joint examination of defense systems and gene flux in *C. difficile*

Until now, we mainly concentrated our attention on the CamA MTase and on its methylation motif CAAAAA. In this section we extended our analysis to the multiple R-M systems associated with the *C. difficile* epigenome and to other defense systems. While the relationship between some defense systems (e.g., CRISPR-Cas and R-M systems) has been studied in an experimental setting[2,3], the rich collection of *C. difficile* methylomes and their genomes provides an unprecedented opportunity to jointly analyze its diversity of defense systems in a data-driven manner.

In addition to R-M systems, we searched for evidence of additional systems: CRISPR-Cas, toxin-antitoxin (T-A), abortive infection (Abi) systems, bacteriophage exclusion (BREX)[4], prokaryotic Argonautes (pAgos)[5], DISARM[6], and a set of 10 recently-discovered defense systems[7] (Supplementary Materials and Methods). Only T-A and CRISPR-Cas systems were ubiquitous in our genomes (Supplementary Figs. 1a-e, Supplementary Tables 3a-d), and all CRISPR-Cas systems detected were of Type-IB[8], consistent with earlier studies[9-11] (Supplementary Fig. 1b).

Different types of defense systems are expected to confer different degrees of protection against invading DNA. Also, under certain conditions, some defense systems may even facilitate genetic exchange between cells, as recently shown for CRISPR-Cas[12] and R-M systems[13]. Thus, we enquired how gene flux was distributed within our dataset (Supplementary Materials and Methods; Supplementary Fig. 1a, Supplementary Table 3e, f), and how is it associated with the multiple defense systems present in it. Specifically, to test the effect of R-Ms, CRISPR-Cas, and T-As (the most abundant defense systems found across *C. difficile* strains) on gene flux, we built stepwise linear models to assess the role of each of these variables in explaining the variance of HGT and HR. Interestingly, we found a strong positive association between CRISPR spacer count and HGT/HR (Supplementary Table 3g, Supplementary Fig. 2). The increase in gene flux with spacer content, although somehow unexpected, could be reconciled if generalized transduction occurs in *C. difficile*[14,15], as recently shown for *Pectobacterium*[12]. R-M and T-A abundance had a less important explanatory role in the prevention of gene flux (Supplementary Table 3g).

Next, we made use of i) the unique information on R-M recognition motifs provided by SMRT-seq, and ii) the exceptional diversity of Type I R-M systems (as well as the near depletion of other types of complete systems) observed in our *C. difficile* dataset, to better understand their impact on phage target side avoidance and gene flux.

Restriction site avoidance is the most effective way to escape the action of R-M systems, and it has been predominantly studied for those belonging to Type II systems[16-19]. To investigate this, we selected representative members of the *Siphoviridae* and *Myoviridae* families and used Markov chain models to compute the number of observed and expected Type I R-M target sites accounting for oligonucleotide composition of each phage genome

(Materials and Methods). We then used as a measure of genetic transfer, the number of recent horizontal gene transfer (HGT) gains from the pattern of presence/absence of gene families in the species tree[20]. Our data suggest that *C. difficile* phages have generally evolved to reduce the number of several Type I recognition sites (Supplementary Fig. 3a). Concomitantly, we found an inverse linear trend between HGT and O/E ratios of Type I motif targets in phages (Spearman's $\rho$ = -0.826, $P < 0.05$) (Supplementary Fig. 3b). This suggests a link between the frequencies at which different *C. difficile* genomes (or STs) are targeted by phages, and the latter's capacity to underrepresent certain motifs targeted by the cell's R-M machinery.

#### Core and pan-genome analysis of *C. difficile*

The core-genome contains 2,118 orthologous protein families (Supplementary Fig. 6b), corresponding to 64.6% of the smallest proteome (*C. difficile* M120). Gene rarefaction analyses showed that the core-genome varies little with the addition of the last genomes (Supplementary Fig. 6b), suggesting that this estimate is robust. *C. difficile* has a large pan-genome with a total of 8,246 gene families (Supplementary Fig. 6b). The spectrum of gene frequencies for the *C. difficile* pan-genome showed that the vast majority of gene families were either encoded in a few genomes (47.9% in four or less) or in most of them (37% in more than 41 genomes) (Supplementary Fig. 6c). Hence, *C. difficile* is characterized by a large and diverse pan-genome and low levels of genome conservation, which fall in the interval of previous estimates[21-23].

#### Inference of homologous recombination events by ClonalFrameML

Homologous recombination (HR) is an evolutionary mechanism that takes place between highly similar sequences[24], and has been heralded as one of the evolutionary forces underlying the emergence of *C. difficile*'s pathogenicity[21,25]. This prompted us to examine whether HR may contribute to the cross-genome variation of CAAAAA motif sites located in the core-genome. We started by quantifying the diversity of gene repertoires (core- and pan-genome) in *C. difficile* (Supplementary Figs. 6b,c; Supplementary Materials and Methods), and then inferred 277 recombination events at the phylogenetic tree tips (Supplementary Table 3e; Supplementary Materials and Methods). While the detailed breakdown of HR events suggests great variation across genomes (Supplementary Fig. 6d), two primary HR peaks were prominent. The first (0.3-0.7 Mb in the core-genome) encompasses genes involved in motility and chemotaxis, such as those belonging to the *flg*, *fli*, and *flh* operons, as well as genes pertaining to ABC and PTS transport systems. The second (2.6-2.7 Mb in the core genome) harbors genes pertaining to the S-layer (*slpA*, *cwp*, etc). Interestingly, both peaks match regions enriched for CAAAAA sites (Fig. 3a), suggesting a possible co-localization. The analysis showed that the recombination to mutation ratio (R/θ) is 0.162, the average length of recombined fragments (δ), is 159.4 bp, and the average distance between donor and recipient is 0.047. Thus, mutations are roughly 6.16 times more frequent than recombination, while the impact of recombination over mutation is 1.22 higher towards the evolution of these strains.

#### Estimation of gene flux in *C. difficile* genomes

We used two measures of genetic transfer: the first based on the number of recent HGT gains and the second based on the number of HR events in the core-genome using two different programs (ClonalFrameML and Geneconv) (Supplementary Materials and Methods). We identified 4,928 events of gene transfer in the *C. difficile* pan-genome (Supplementary Table 3e, Supplementary Fig. 1a). These events were very unevenly distributed across branches (and STs), from no events in CD_020265, to 530 in

CD_020486. For HR, we found 403 and 270 events in the core-genome as given by Geneconv and CFML respectively (Supplementary Table 3e, Supplementary Fig. 1a). Even if the two programs provided different numbers of events, their results per genome were significantly correlated (Spearman's correlation = 0.60, $P < 10^{-3}$). To further complement the information on HR and HGT, we performed a one-way hierarchical clustering with the number of matches between spacers and currently known *Clostridium* phage sequences (Supplementary Materials and Methods). We found very heterogeneous targeting profiles across isolates (Supplementary Fig. 1a, Supplementary Table 3f), which still correlated with HGT and HR (both with Spearman's correlation = 0.33, $P < 0.05$).

### Supplementary Materials and Methods

#### Presence and conservation of *camA* in *C. difficile* isolates

To investigate the pervasive role and conservation of *camA*, we searched for its presence in a global and UK collection of *C. difficile* 027/BI/NAP1 (n=291)[26] genomes (Supplementary Table 4). For this, SRA Illumina reads were converted to FASTQ files using fastq-dump v2.8.0 and subsequently mapped to the *C. difficile* 630 reference genome using Bowtie2 v2.2.9[27] in paired-end mode. The resulting SAM files were converted to BAM format (with removal of unmapped reads and PCR duplicates), and sorted using SAMTOOLS v1.3.1[28]. To assess coverage, sequence depths were computed using the genomeCov function of BEDTOOLS v2.26.0 for each strand separately. Variant sites were called from the aligned reads using the *mpileup* and *bcftools* tools in SAMTOOLS.

#### Identification of defense systems

CRISPR repeats were identified using the CRISPR Recognition Tool (CRT) v1.2[29] with default parameters. For CRISPR spacer homology search, we considered as positive hits those with at least 80% identity. For *cas* gene identification, we obtained Cas protein family HMMs from the TIGRFAM database[30] v15.0 and PFAM families annotated as Cas families (downloaded from ftp://ftp.ncbi.nih.gov/pub/wolf/_suppl/CRISPRclass/crisprPro.html). In total we collected 129 known Cas protein families (98 TIGRFAMS and 31 PFAMs), which were used for similarity searching. Genes pertaining to abortive infection (Abi) systems were searched with the PFAM profiles PF07751, PF08843, and PF14253 (last accessed in January 2018). Bacteriophage Exclusion (BREX) systems were searched using PFAM profiles for the core genes *pglZ* (PF08655) and *brxC*/*pglY* (PF10923), and specific PFAM profiles for each BREX type as indicated previously[4]. DISARM systems were identified using the PFAM signature domains (PF09369, PF00271, PF13091) belonging to the core gene triplet characteristic of this system[6]. To search for prokaryotic Argonaute (pAgo) genes we built a dedicated HMM profile based on a list of 90 Ago-PIWI proteins[31]. Searches for the ensemble of newly found antiphage systems were performed using the list of PFAM profiles published by the authors[7]. Type II toxin-antitoxin (T-A) systems were detected using the TAFinder tool[32] with default parameters. Matches of CRISPR spaces were performed against well-known *C. difficile* phages: five siphophages (ϕCD111 (NC_028905.1), ϕCD146 (NC_028958.1), ϕCD38-2 (NC_015568.1), ϕCD6356 (NC_015262.1), ϕCD211 (NC_029048.2)), five small-tail myophages (ϕMMP04 (NC_019422.1), ϕCD506 (NC_028838.1), ϕCDHM11 (NC_029001.1), ϕCD481-1 (NC_028951.1), ϕCDHM13 (NC_029116.1)), five medium-tail myophages (ϕMMP03 (NC_028959.1), ϕCDMH1 (NC_024144.1), ϕC2 (NC_009231.1), ϕCD119 (NC_007917.1), ϕCDHM19 (NC_028996.1)), and four long-tail myophages (ϕCD27 (NC_011398.1), ϕMMP02 (NC_019421.1), ϕCD505 (NC_028764.1), ϕMMP01 (NC_028883.1)).

#### Identification and classification of prophages, conjugative/mobilizable elements and integrons

Prophages were detected using Phage Finder v2.1[33] under strict mode, and PHASTER[34] under default settings. We took the common hits obtained by both programs, as well as those very few cases (~10% of the hit list) corresponding to complete prophages predicted by just one of the programs. All elements smaller than 18 kb, or lacking matches to core phage proteins (e.g. terminase, capsid, head, tail proteins) were removed (Supplementary Table 2c). Integrons were searched with IntegronFinder[35] under default settings. The identification of genes encoding the functions related to conjugation in integrative conjugative elements (ICEs) was performed as previously described[36]. Briefly, an element was considered as conjugative when it contained the following components of the conjugative system: a VirB4/TraU ATPase, a relaxase, a coupling ATPase (T4CP), and a minimum number of mating pair formation (MPF) type-specific genes: two for types $MPF_{FA}$ and $MPF_{FATA}$, or three for the others (types F, T, and G). In the case of integrative mobilizable elements (IMEs), they were identified by the fact that they encode relaxases but lack a complete conjugative transfer system, which is encoded in *trans* by another mobile element. Delimitation of ICEs and IMEs was performed considering flanking core genes as upper bounds for their extremities.

#### Phylogenetic analyses

The reference phylogenetic tree of *C. difficile* was built from the concatenated alignment of protein families of the core-genome using MUSCLE v3.8.31 (default parameters). Since at this evolutionary distance the DNA sequences provide more phylogenetic signal than protein sequences, we back-translated the alignments to DNA. Poorly aligned regions were removed with BMGE[37]. The tree was computed with RAxML[38] v8.00 under the GTR model and a gamma correction (GAMMA) for variable evolutionary rates. 100 bootstraps were performed on the concatenated alignment to assess the robustness of the topology of the tree.

#### Bacterial strains and growth conditions

The 630Δ*erm*Δ*pyrE* parental strain was used for *pyrE*-based allelic-coupled exchange (ACE[39]). See Supplementary Table 5a for a list of *C. difficile* and *E. coli* strains. *C. difficile* strains were grown from frozen stocks on brain heart infusion media (BHIS) plates supplemented with taurocholate (TA, 0.1% w/v; 1.9 mM), kanamycin (50 µg/mL), and cefoxitin (8 µg/mL) as needed. For ACE, *C. difficile* defined media (CDDM)[40] was supplemented with 5-fluoroorotic acid (5-FOA) at 2 mg/mL and uracil at 5 µg/mL. Cultures were grown at 37 °C under anaerobic conditions using a gas mixture containing 85% $N_2$, 5% $CO_2$, and 10% $H_2$. The growth curves were performed in BHIS media with gentle shaking. *E. coli* strains were grown at 37 °C, shaking at 225 rpm in Luria-Bertani broth (LB). The media was supplemented with chloramphenicol (20 µg/mL) and ampicillin (50 µg/mL) as needed.

#### *E. coli* strain construction

Primers used in this manuscript are listed in Supplementary Table 5b. *C. difficile* 630 genomic DNA was used as the template. To clone the pMTL-YN3-Δ*camA* construct, primer pairs #2332 and 2334 and #2333 and 2335 were used to amplify the region 662 bp upstream and 226 bp downstream of *CD630_27580,* respectively. The resulting PCR products were cloned into pMTL-YN3 using Gibson assembly[41]. This construct encodes a *CD630_27580* deletion in which the first 14 codons are linked to the last 139 codons with an intervening stop codon between the 5' and 3' end of the gene to avoid production of the last 139 amino acids of CamA. To clone the *camA* complementation constructs, primer pair #2286 and 2287 was used to amplify *camA* and 163 bp of its upstream region. The resulting PCR product was recombined into pMTL-YN1C by Gibson assembly. The *N165A* complementation construct was cloned in a similar fashion except that the primer pairs consisted of #2286 and #2532 and #2531 and #2287. The plasmids were transformed into *E. coli* DH5α, and the resulting plasmids were confirmed by sequencing using Genewiz and then transformed into HB101/pRK24 for conjugations.

#### *C. difficile* strain construction

ACE was used to construct 630Δ*erm*Δ*pyrE*Δ*camA* using uracil and 5-fluoroorotic acid to select for plasmid excision as previously described[42]. The flanking primer pair #2274 and #2279 was used to screen for the *camA* deletion as shown in Supplementary Fig. 5a (primers are provided in Supplementary Table 5b). Colonies that appeared to harbor gene deletions were validated by performing an internal PCR using a primer (#2288) that binds within the region deleted and a primer (#2279) that binds to the region flanking the deletion. Two independent clones from the allelic exchange were phenotypically characterized. The *camA* complementation strains were constructed as previously described by using CDDM plates to select for restoration of the *pyrE* locus via recombination[42]. Two independent clones from each complementation strain were phenotypically characterized.

#### Cell length measurements

Cells were grown to mid-log and stationary phase in BHIS broth or sporulation was induced as described below for three biological replicates. Cells were imaged by phase contrast microscopy on a Zeiss Axioskop with a 100x Zeiss Plan-Neo-fluar objective (1.3 NA) at each timepoint. Cell length was calculated using the MicrobeJ plugin for Fiji/ImageJ[43]. Image thresholding was done using the local default method in MicrobeJ/Fiji to account for variations in background. Cell detection parameters were optimized (Area: 0-20 μm$^2$, Length: 1 μm-max, Width: 0.5-1 μm) and contours were generated using an interpolated rod-shaped method. Cell length data was exported from MicrobeJ and analyzed using Prism 8 (Graph-pad).

#### Biofilm assays

Biofilm assays were done as previously described[44]. Briefly, overnight cultures of *C. difficile* were diluted 1:100 in BHIS-1% glucose-50 mM sodium phosphate buffer (pH 7.5) in 24-well polystyrene plates. After 24 hours of growth at 37℃, supernatants were removed, the biofilms were washed once with PBS and then stained for 30 minutes with 0.1% (w/v) crystal violet. After 30 minutes, the biofilms were washed again with PBS, and the crystal violet was solubilized with ethanol. Absorbance was read at 570 nm. Three independent experiments were performed, with each strain assayed in quadruplicate in each experiment.

#### Sporulation

*C. difficile* strains were inoculated from glycerol stocks overnight onto BHIS-TA plates. Liquid BHIS cultures were inoculated from colonies arising on these plates. The cultures were grown to early stationary phase, back-diluted 1:50 into BHIS, grown until they reached an $OD_{600}$ between 0.35 and 0.75, and then 120 µL of this culture was spread onto 70:30 plates (40 mL). Sporulating cultures were harvested into phosphate-buffered saline (PBS), the sample was pelleted, and sporulation levels were visualized by phase-contrast microscopy as previously described[45].

#### Fluorescence microscopy

Fluorescence microscopy was performed on sporulating cultures using Hoechst 33342 (Molecular Probes; 15 µg/ml) and FM4-64 (Invitrogen; 1 µg/ml) to stain nucleoid and membrane, respectively. Cells were mounted on a 1% agarose in PBS pad. Images were acquired on a Nikon 80i upright epifluorescence microscope using a Nikon 60x plan apochromat phase contrast objective (1.4 NA) in 12-bit format using Nikon NIS elements software. Images were processed in Adobe Photoshop CC for adjustment of brightness, contrast levels, and pseudocoloring.

#### Spore purification

Sporulation was induced on four 70:30 plates for 48-65 h for each strain tested as described above, and spores were purified as previously described[46]. Briefly, sporulating cultures were scraped up, washed repeatedly in ice-cold water, incubated overnight in water on ice, treated with DNase I (New England Biolabs) at 37 ˚C for 45-60 min, then purified on a density gradient (Histodenz, Sigma Aldrich). Spores were resuspended in 600 µL water for final storage at 4 ˚C. Spore purity was assessed using phase contrast microscopy (>95% pure), and the optical density at 600 nm was measured. Spore purification yields were determined from three independent spore preparations. Statistical significance was determined using a one-way ANOVA and Tukey's test.

#### Heat resistance assay

Heat-resistant spore formation was measured in sporulating *C. difficile* cultures after 20-24 h as previously described[45]. The heat resistance ($H_{RES}$) efficiency represents the average ratio of heat-resistant colony forming units (CFUs) to total CFUs for a given strain relative to the average ratio determined for wild type. $H_{RES}$ was determined based on the average $H_{RES}$ values for a given strain in three biological replicates. Statistical significance was determined using a one-way ANOVA and Tukey's test.

#### Germination assay

Germination assays were performed as previously described[47]. Spores (0.35 $OD_{600}$ units, corresponding to ~1 x $10^7$) were resuspended in 100 µl of water, and 10 µL of this mixture was removed for 10-fold serial dilutions in PBS. The dilutions were plated on BHIS-TA, and colonies arising from germinated spores were enumerated after 18-21 h. Germination efficiencies were calculated by averaging the CFUs produced by spores for a given strain relative to the number produced by wild-type spores for three biological replicates. Statistical significance was determined by performing a one-way ANOVA on natural log-transformed data using Tukey's test. The data were transformed because the use of independent spore preparations resulted in a non-normal distribution. Regardless, no statistical significance in germination efficiency was observed for the mutant and its complements.

#### Spore chloroform resistance

Spores (0.75 $OD_{600}$ units, corresponding to ~2 x $10^7$ spores) were re-suspended in 190 µL water. 90 µL were then added to tubes containing either 10 µL of water or chloroform for 15 min after which 10 µL of the sample was serially diluted in PBS and plated on BHIS-TA as described previously[46,48].

#### Animal infection studies

All animal experimentation was performed under the guidance of veterinarians and trained animal technicians within the University of North Carolina Division of Comparative Medicine. Animal experiments were performed with prior approval from the UNC Institutional Animal Care and Use Committee. Animals considered moribund as defined in the protocols were euthanized by $CO_2$ asphyxiation followed by a secondary, physical method in accordance with the Panel on Euthanasia of the American Veterinary Medical Association. The University complies with state and federal Animal Welfare Acts, the standards and policies of the Public Health Service.

**Murine model**: The parental *C. difficile* strain 630Δ*erm*, the MTase mutant 630Δ*erm*Δ*camA*, and the MTase complemented strain were evaluated in an antibiotic-treated mouse model as previously described[49,50]. Groups of 8- to 10-week old female and male (N = 6 each) C57BL/6 mice (*Mus musculus*; Charles River Laboratories) were administered a cocktail of antibiotics (kanamycin (400 μg/ml), gentamicin (35 μg/ml), colistin (850 units/ml), vancomycin (45 μg/ml), and metronidazole (215 μg/ml)) in their water *ad libitum* seven days prior to inoculation for three days, followed by a single intra-peritoneal dose of clindamycin (10 mg/kg body weight) 2 days prior to inoculation. Mice were inoculated with $10^5$ spores by oral gavage. Mock-inoculated animals were included as controls. Cage changes were performed every 48 h post-inoculation. Fecal samples were collected every 24 h for seven days post-inoculation. Dilutions were plated on BHIS-agar containing 0.1% of the germinant taurocholate to enumerate spores as colony forming units (CFU) per gram of feces.

**Hamster model**: The above strains were tested in male and female (N = 3 each) Syrian golden hamsters strain LVG (*Mesocricetus auratus*; Charles River Laboratories) as described previously[51]. Hamsters were administered a single dose of clindamycin (30 mg/kg body weight) by oral gavage, then inoculated with approximately 5,000 spores of the above strains 5 days later. Mock-inoculated animals were included as controls. Hamsters were monitored for weight loss and diarrheal symptoms and were considered moribund after 15-20% weight loss from maximum body weight, with or without concurrent diarrhea.

#### RNA processing

For analyses of sporulating cell transcriptomes, RNA was extracted from three biological replicates of wild type and Δ*camA* growing on 70:30 sporulation media after 9 and 10.5 h of growth using the FastRNA Pro Blue Kit (MP Biomedical) and the FastPrep-24 automated homogenizer (MP Biomedical), similar to previous work[52]. For analyses of the mid-log and early stationary phase cultures, overnight cultures of wild type and Δ*camA* in BHIS were back-diluted 1:50 into three biological replicates of 30 mL of BHIS in 125 mL Erlenmeyer flasks. The cultures were grown until mid-log phase ($OD_{600}$ = 0.5-0.6) and early stationary phase ($OD_{600}$ = 1.3-1.4). RNA was harvested from 15 mL and 10 mL of the same cultures for the mid-log and early stationary phase cultures, respectively. Contaminating genomic DNA was depleted using three successive DNase treatments, with the last treatment being on column using the Qiagen RNeasy kit. Samples were tested for genomic DNA contamination using quantitative PCR for 16S rRNA and the *sleC* gene. DNase-treated RNA (15 µg) was enriched for mRNA using the Ribo-Zero Magnetic Kit (Epicentre) for the broth-grown cultures. Ribosomal RNA was depleted from RNA harvested from sporulating cultures using the Ambion MICROB*Express* Bacterial mRNA Enrichment Kit (Thermo Fisher) because Ribo-Zero kits were temporarily discontinued. The quality of total RNA was validated using an Agilent 2100 Bioanalyzer. Samples for qRT-PCR analyses were harvested in triplicate from a separate set of three biological replicates that were grown identically to the cultures used for RNA-Seq analyses. The RNA was processed similarly except that mRNA enrichment was done using a MICROB*Express*, and the DNase-treated RNA samples for qRT-PCR analyses were tested for genomic DNA contamination using quantitative PCR for *rpoB*.

#### Quantitative real-time PCR (qRT-PCR)

Transcript levels were determined from cDNA templates prepared from the three biological replicates described above. Gene-specific primer pairs are provided in Supplementary Table 5b. qRT-PCR was performed as described[53], with the exception that we have used an iTaq Universal SYBR Green supermix (BioRad), 50 nM of gene specific primers and a Mx3005P qPCR system (Stratagene) in a total volume of 25 µl. The following cycling conditions were used: 95 °C for 2 min, followed by 40 cycles of 95 °C for 15 s and 60 °C for 1 min. Transcript levels were normalized to the housekeeping gene *rpoB* using the standard curve method.

#### Western blots

**Sporulation protein analyses**: Sporulation was induced as indicated, and samples were harvested and processed for immunoblotting as previously[46]. Total protein in each sample was quantified using the Pierce 660nm protein assay with the ionic detergent compatibility reagent (Thermo Fisher) and 5 µg of protein was loaded for each sample. $\sigma^F$, $\sigma^E$, and Spo0A were resolved on 15% SDS-PAGE gels while SpoIIQ and SpoIVA were resolved on 12% SDS-PAGE gels. Protein was transferred to PVDF membranes, which were subsequently probed with rabbit ($\sigma^F$, $\sigma^E$, SpoIIQ) and mouse (Spo0A, SpoIVA) polyclonal primary antibodies and α-rabbit IR800/α-mouse IR680 secondary antibodies (LI-COR). Blots were imaged on the LiCor Odyssey CLx. Results shown are representative of analyses of three biological replicates.

**Toxin analyses**: Overnight cultures of *C. difficile* were diluted 1:50 in TY medium and incubated at 37℃ for 24 h. Cells were collected by centrifugation, suspended in SDS-PAGE buffer, and boiled for 10 min. Samples were then run on 4–20% Mini-PROTEAN TGX Precast Protein Gels (Bio Rad) and transferred to a nitrocellulose membrane. TcdA was detected as described previously using a mouse α-TcdA primary antibody (Novus Biologicals) and goat anti-mouse IgG conjugated with IR800 (Thermo Fisher)[54].

#### Identification of the core- and pan-genome

The *C. difficile* core-genome was built using a methodology previously published[55]. Briefly, a preliminary list of orthologs was identified as reciprocal best hits using end-gap-free global alignment between the proteome of a pivot (*C. difficile* 630) and each of the other strain's proteomes. Hits with <80% similarity in amino-acid sequence or >20% difference in protein length were discarded. This list of orthologs was then refined for every pairwise comparison using information on the conservation of gene neighborhood. Positional orthologs were defined as bi-directional best hits adjacent to at least four other pairs of bi-directional best hits within a neighborhood of 10 genes (five upstream and five downstream). The core-genome of each clade was defined as the intersection of pairwise lists of positional orthologs. The pan-genome was built using the complete gene repertoire of *C. difficile*. We determined a preliminary list of putative homologous proteins between pairs of genomes by searching for sequence similarity between each pair of proteins with BLASTP (default parameters). We then used the *e*-values ($<10^{-4}$) of the BLASTP output to cluster them using SILIX (v1.2.11, http://lbbe.univ-lyon1.fr/SiLiX)[56]. We set the parameters of SILIX such that two proteins were clustered in the same family if the alignment had at least 80% identity and covered >80% of the smallest protein (options –I 0.8 and –r 0.8). Core- and pan-genome accumulation curves were built using a dedicated R script. Regression analysis for the pan-genome was performed as described previously[57] by the Heap's power law $n = k \cdot N^{-\alpha}$, where *n* is the pan genome family size, *N* is the number of genomes, and *k*, $\gamma$ ($\alpha = 1 - \delta$) are specific fitting constants. For $\alpha > 1$ ($\delta < 0$) the pan-genome is considered closed, i.e. sampling more genomes will not affect its size. For $\alpha < 1$ ($0 < \delta < 1$) the pan-genome remains open and addition of more genomes will increase its size.

#### Inference of homologous recombination

We inferred homologous recombination on the multiple alignments of the core-genome of *C. difficile* (ordered LCBs obtained by progressiveMauve were used) using

ClonalFrameML v10.7.5[58] and Geneconv v1.81a[59]. The first used a predefined tree (*i.e.* the clade's tree), default priors $R/\theta = 10^{-1}$ (ratio of recombination and mutation rates), $1/\delta = 10^{-3}$ (inverse of the mean length of recombination events), and $\nu = 10^{-1}$ (average distance between events), and 100 pseudo-bootstrap replicates, as previously suggested[58]. Mean patristic branch lengths were computed with the R package "ape" v3.3[60], and transition/transversion ratios were computed with the R package "PopGenome" v2.1.6[61]. The priors estimated by this mode were used as initialization values to rerun ClonalFrameML under the "per-branch model" mode with a branch dispersion parameter of 0.1. The relative effect of recombination to mutation (r/m) was calculated as $r/m=R/\theta\times\delta\times\nu$. Geneconv was used with options /w123 to initialize the program's internal random number generator and –Skip_indels which ignores all sites with missing data.

#### Reconstruction of the evolution of gene repertoires

We assessed the dynamics of gene family repertoires using Count[20] (downloaded in January 2018). This program uses birth-death models to identify the rates of gene deletion, duplication, and loss in each branch of a phylogenetic tree. We used presence/absence pan-genome matrix and the phylogenetic birth-and-death model of Count, to evaluate the most likely scenario for the evolution of a given gene family on the clade's tree. Rates were computed with default parameters, assuming a Poisson distribution for the family size at the tree root and uniform duplication rates. One hundred rounds of rate optimization were computed with a convergence threshold of $10^{-3}$. After optimization of the branch-specific parameters of the model, we performed ancestral reconstructions by computing the branch-specific posterior probabilities of evolutionary events, and inferred the gains in the terminal branches of the tree. The posterior probability matrix was converted into a binary matrix of presence/absence of HGT genes using a threshold probability of gain higher than 0.2 at the terminal branches. To control for the effects of the choices made in the definition of our model, we computed the gain/loss scenarios using the Wagner parsimony (same parameters, relative penalty of gain with respect to loss of 1). The HGT events inferred by maximum likelihood and those obtained under Wagner's parsimony were highly correlated (Spearman's $\rho = 0.96$, $P < 10^{-4}$).

### Supplementary Figures

**Supplementary Fig. 1.** Multiple defense systems and gene flux control in *C. difficile*. (a) Heatmap aggregate depicts: abundance of defense systems (R-M, abortive infection (Abi), average number of spacers per CRISPR, toxin-antitoxin (T-A), and Shedu systems (other)), homologous recombination (HR) events (given by Geneconv and ClonalFrameML (CFML)), horizontal gene transfer (HGT, given by Wagner parsimony), and number of phage-targeting CRISPR spacers (Supplementary text). Phages were clustered according to their family (*Siphoviridae* (S), *Myoviridae* (M)), and tail type. (b) Cas genes detected in *C. difficile*. Apart from the complete Type-IB gene cluster (*cas1*-*cas8*), we also observed two truncated gene clusters lacking *cas1*, *cas2*, and *cas4*. One of the truncated operons was present across all genomes, while the second was restricted to ST-1 and ST-55. (c) Example of a putative 'defense island' detected in CD_020472 harboring: a Druantia-like system, two T-A systems, two solitary MTases, and one Type I R-M system. The Druantia-like system is similar to the previously reported Type II Druantia systems[7] in the sense that a PF00271 helicase conserved C-terminal domain and DUF1998 (PF09369) are associated with a nearby cytosine methylase. However, it lacks a PF00270 DEAx box helicase. (d) Genomic context of the *sduA* gene in CD_22456 pertaining to the newly identified Shedu defense system. The gene is located in an integrative conjugative element (ICE) (Supplementary Table 2d). (e) Observed/expected (O/E) ratios for co-localized defense systems (maximum of 10 genes apart). Only the most abundant systems were included in the analysis. Expected values were obtained by multiplying the total number of defense systems by the fraction of co-localized defense systems.

**Supplementary Fig. 2.** Relation between gene flux and CRISPR spacer content. (a) Association between genetic flux (horizontal gene transfer (HGT) and homologous recombination (HR, computed using both ClonalFrameML (CFML) and Geneconv)) and number of CRISPR spacers. The latter were used as proxy of their activity. Data was plotted after excluding very similar ST-1 genomes. The criteria to remove these genomes were based on similarities in R-M content, and gene flux, i.e., all ST-1 genomes but CD_020475, CD_020474, CD_021026 were removed. (b) Same as (a) but considering the complete dataset.

**Supplementary Fig. 3.** Interplay between Type I R-M systems and gene flux in *C. difficile*. (a) Observed/expected (O/E) ratios for Type I target recognition motifs in *Clostridium* phage genomes. 6 phage genomes representative of *Siphoviridae* and *Myoviridae* families and tail types were analyzed (ϕCD111, ϕCDHM11, ϕMMP01, ϕMMP04, ϕC2, ϕCD38). For each motif, we tested if the median value of the O/E ratio in phage genomes was significantly different from 1 with the Mann-Whitney test. O/E values were obtained with R'MES using Markov chain models that take into consideration oligonucleotide composition (Supplementary Materials and Methods). (b) Relation between HGT and O/E ratio for Type I target recognition motifs. For those *C. difficile* genomes harboring a single Type I R-M system (i.e., without the confounding effect of multiple systems), we computed the average values of HGT, and plotted these values against the average O/E ratio for the corresponding target recognition motif in phage genomes. This was only possible for the motifs indicated in brackets. * $P<0.05$, ** $P<10^{-2}$, *** $P<10^{-3}$, Mann-Whitney-Wilcoxon test.

**Supplementary Fig. 4.** Genomic context and conservation of *camA*. (a) CamA protein alignment among *Clostridiales* (*C. mangenotti* LM2 (587 aa, 56% identity), *C. sordellii* (598 aa, 53% identity), *C. bifermentans* WYM (579 aa, 53% identity), *C. dakarense sp. nov* (580 aa, 63% identity), *Peptostreptococcaceae bacterium* VA2) and *Fusobacteriales* (*Psychrilyobacter atlanticus* DSM 19335) using ClustalX. The nine conserved motifs (I-VIII and X) typically found in MTases are highlighted. (b) Phylogenetic tree obtained from the MTase alignment. (c) Phylogenetic tree of the 36 *C. difficile* strains colored by clade (hypervirulent, human/animal (HA) associated) and MLST sequence type (ST). Shown is the genomic context of *camA* across the entire dataset. (d) Expanded view of the region shown in Fig. 1f. The example shown (including coordinates) refers to the reference genome of *C. difficile* 630. + and – signs correspond to the sense and antisense strands respectively. Vertical bars correspond to the distribution of the CAAAAA motif.

**Supplementary Fig. 5.** ∆*camA* construction, purified spore analyses, broth culture growth, and sporulation kinetics. (a) PCR to distinguish between wild-type *camA* and ∆*camA* using flanking primers and primers internal to the deletion. (b) Growth curves comparing wild-type *camA,* ∆*camA*, ∆*camA-C*, and *camA/N165A* cultures grown in BHIS media shaking over time. Early stationary-phase cultures were diluted to a starting O.D. of 0.05 in BHIS media and growth was measured over 9 h. (c) Phase-contrast microscopy analyses of sporulating culture samples prior to and after spore purification on a density gradient. No gross differences in spore morphology were observed between wild type and the MTase mutant. The germination efficiency (G.E.) of purified spores from the indicated strains is shown below. Scale bar represents 5 µm. (d) Chloroform resistance of purified ∆*camA* spores relative to wild type. Spores were treated with 10 % chloroform for 15 min after which spore viability was measured by plating untreated and chloroform-treated spores on media containing germinant and measuring colony forming units. No significant differences in germination efficiency or chloroform resistance were observed. (e) Heat-resistance ($H_{RES}$) efficiencies of sporulating cultures 22 h after sporulation induction were determined relative to wild-type. Three replicates per group were used. Statistical analyses were performed with a one-way ANOVA with Tukey's test. *** $P < 10^{-3}$.

**Supplementary Fig. 6.** Core- / pan-genome analyses of *C. difficile* and HR landscape. (a) Observed/expected (O/E) ratios of the CAAAAA motif in the *C. difficile* chromosome, intragenic, extragenic, and regulatory regions (defined as the windows spanning 100 bp upstream the start codon to 50 bp downstream). Expected values were computed by R'MES using Markov models that take sequence composition into consideration (the highest possible order was used (corresponding to a *k*-mer size of 5)). (b) Core- and pan-genome sizes of *C. difficile*. The pan- and core-genomes were used to perform gene accumulation curves. These curves describe the number of new genes (pan-genome) and genes in common (core-genome) obtained by adding a new genome to a previous set. The procedure was repeated 1,000 times by randomly modifying the order of integration of genomes in the analysis. The values for the specific constants obtained after Heap's law fitting (Supplementary Materials and Methods) are 2,887 and 0.271, respectively for the $k$ and $\gamma$, thus implying an open pan-genome. (c) Spectrum of frequencies for *C. difficile* gene repertoires. It represents the number of genomes where the families of the pan-genome can be found, from 1 for strain-specific genes to 45 for core-genes. Red indicates accessory genes and blue the genes that are highly persistent in *C. difficile.* (d) Graphical representation of the recombinational events as inferred by ClonalFrameML. The HA and hypervirulent branches of the tree are depicted in colors. Substitutions are represented by vertical lines and recombination events by dark blue horizontal bars. Light blue vertical lines represent the absence of substitutions, and white lines refer to non-homoplasic substitutions. All other colors represent homoplasic substitutions, with increases in homoplasy associated with increases in the degree of redness (from white to red). (e) O/E ratios of orthologous variable CAAAAA motifs in the core-genome (excluding recombination tracts) versus recombination tracts and (f) core versus accessory genome. *P*-values correspond to the Chi-square test.

**Supplementary Fig. 7.** Non-methylated CAAAAA motif sites. Motif overlap with TFBS and TSS. (a) Interpulse duration ratio (ipdR) density distribution of the terminal adenine of CAAAAA. Motifs were considered as non-methylated if the terminal adenine had IPD ratios <1.5 (stippled line), coverage was >20×, and methylation scores <20 (gray distribution). Also shown for comparison are the sections delimited by quantiles (Q) 1, 5, 10, and 50. (b) Additional examples of highly conserved non-methylated CAAAAA motif sites (red ovals) and corresponding genetic context. Positions indicated above the graph correspond to the non-methylated base. (c) %CAAAAA motif sites (non-methylated (NM) and methylated (M)) overlapping CodY and XylR TFBS for each *C. difficile* isolate excluding the 13 genomes belonging to ST1. (d) Additional examples of chromosomal regions for which non-methylated CAAAAA motif sites overlap TSSs (shown as arrows). (e) %CAAAAA motif sites (non-methylated and methylated) overlapping TSSs for each *C. difficile* isolate (excluding genomes belonging to ST1). (f) % mutated reads (SNPs + indels) in CAAAAA, GAAAAA, and TAAAAA motifs for *M. tuberculosis* (MT), *E. coli* (EC), *C. difficile* (CD) and *H. pylori* (HP). * $P < 0.05$, *** $P < 10^{-3}$, Mann-Whitney-Wilcoxon test.

**Supplementary Fig. 8.** Principal Component Analysis (PCA) and MA-plots for RNA-seq data. (a) PCA performed using DESeq2 rlog-normalized RNA-seq data. (b) MA-plots showing the variation of fold change with mean normalized counts (MNC). Red-colored points have adjusted *P* values < 0.1. Points that fall out of the window are plotted as open triangles pointing either up or down.

**Supplementary Fig. 9**. DE, gene, and protein expression analyses. (a) Enrichment of the CAAAAA motif in DE genes compared to non-DE ones either globally (left) or at each time point studied (right). (b) Time-course change in the expression of genes under the control of the specific sigma factors ($\sigma^F$, $\sigma^E$, $\sigma^G$, and $\sigma^K$) and master transcriptional activator Spo0A. (c) Representative immunoblot time-course comparing the levels of the early sporulation proteins $\sigma^F$, SpoIIQ, $\sigma^E$, and SpoIVA in WT and Δ*camA* at 8, 10, 12, 14, and 16 h following induction of sporulation. (d) Western blot for TcdA for the four *C. difficile* different strains. (e) qRT-PCR of *spoVD* and *cwp17* genes from exponential and stationary phase liquid broth cultures. * $P < 0.05$, ** $P < 10^{-2}$, *** $P < 10^{-3}$.

**Supplementary Fig. 10.** Overlap between multiple datasets of DE genes. Comparisons were performed between DE genes called in this study for each time point (blue-shaded) and those obtained from (a) Jenior *et al*. and (b) Fletcher *et al*. (green-shaded). The dataset of DE genes computed from the study of Jenior *et al*. correspond to a comparison between growths under antibiotic-treated conditions versus growth in a germfree control murine model[62]. The dataset of DE genes computed from the study of Fletcher *et al*. were obtained from murine gut isolates at increasing time points after infection[63]. Color intensities of the outermost layer represent the *P*-value significance of the intersections (3,896 genes were used as background). The height of the corresponding bars is proportional to the number of common genes in the intersection (indicated at the top of the bars for pairwise comparisons between the different studies).
