## Supplementary figures and images for "Epigenomic and functional characterization of a core DNA methyltransferase in the human pathogen *Clostridium difficile*"

### Supplementary Figures

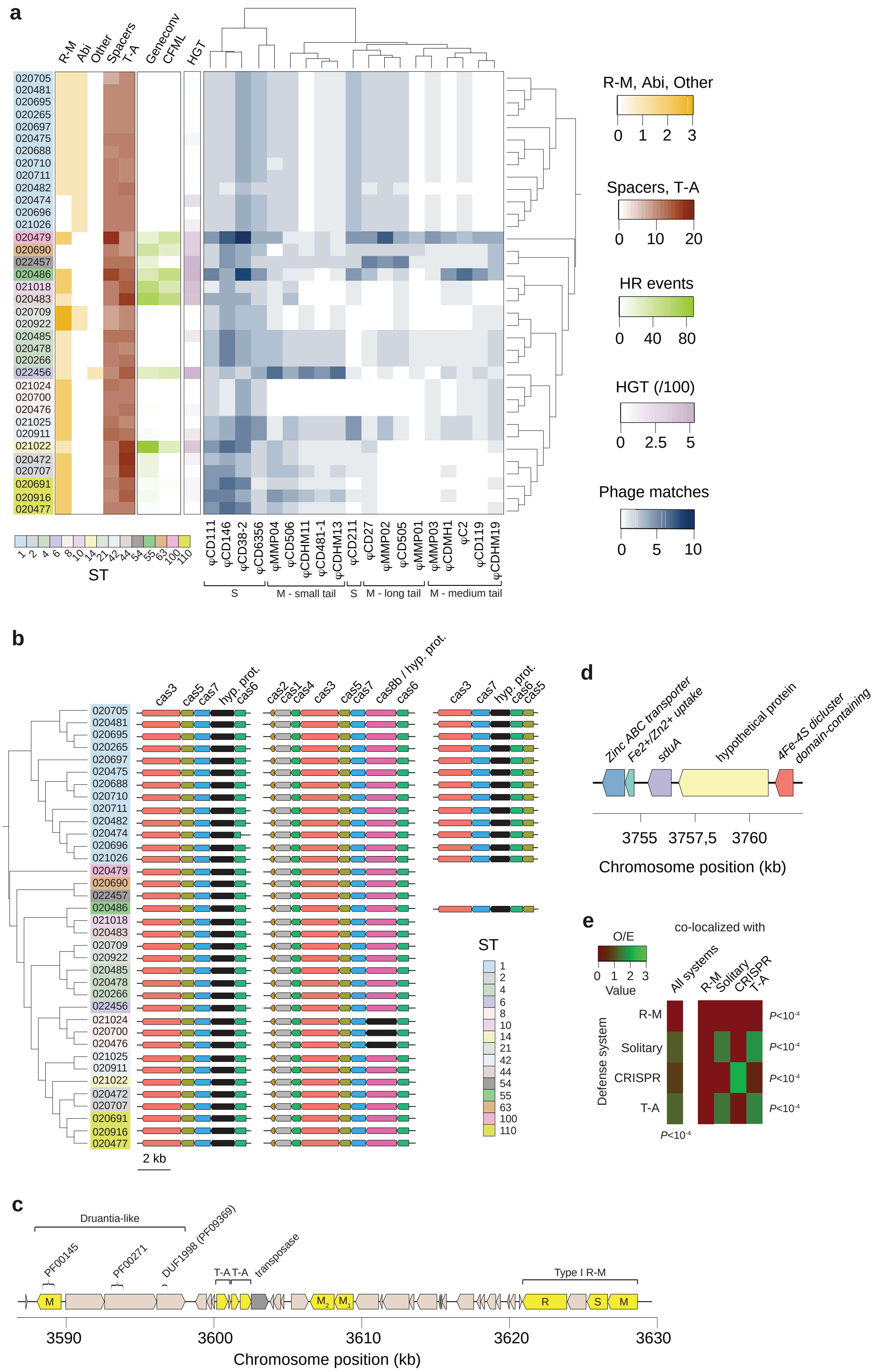

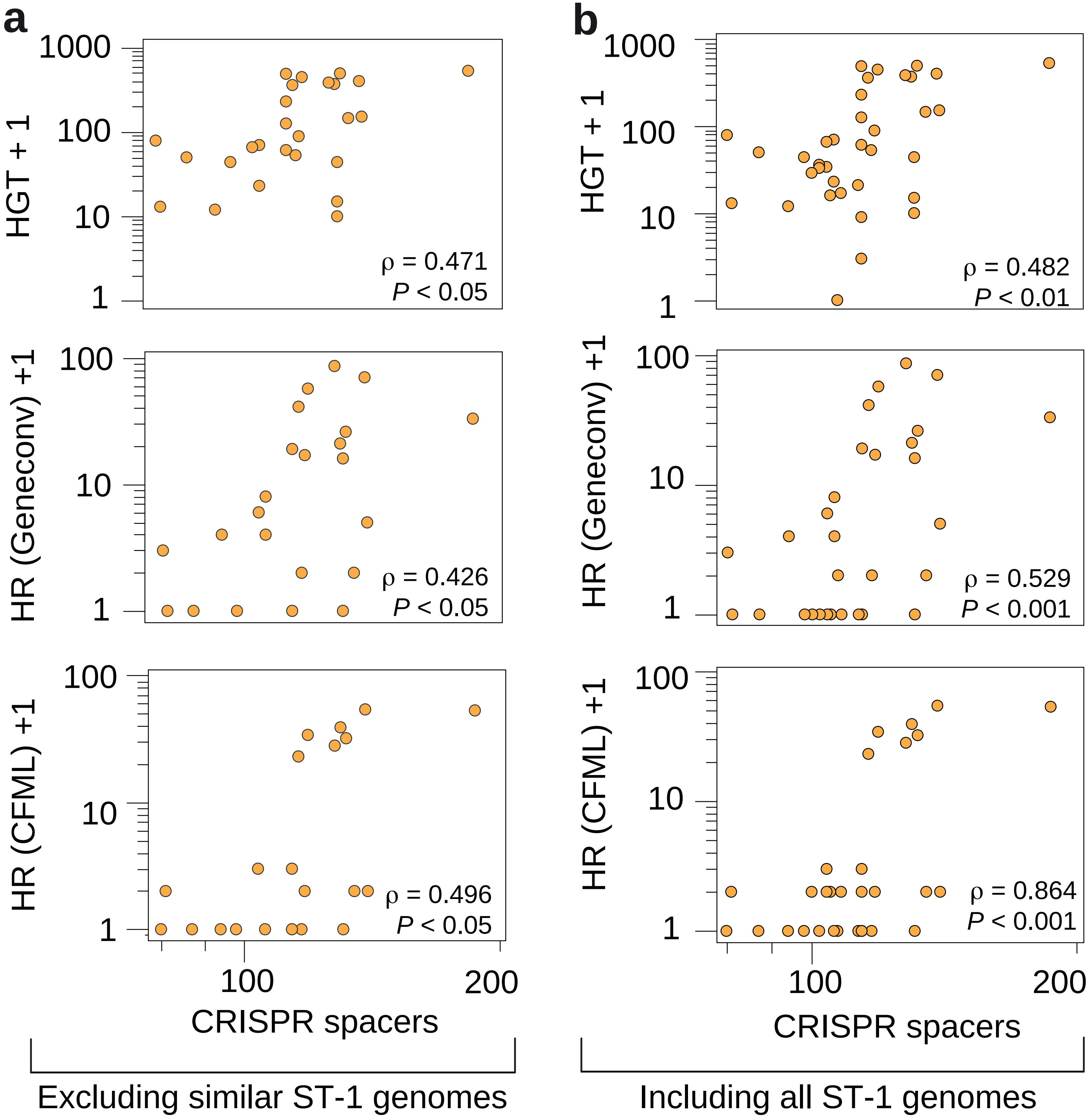

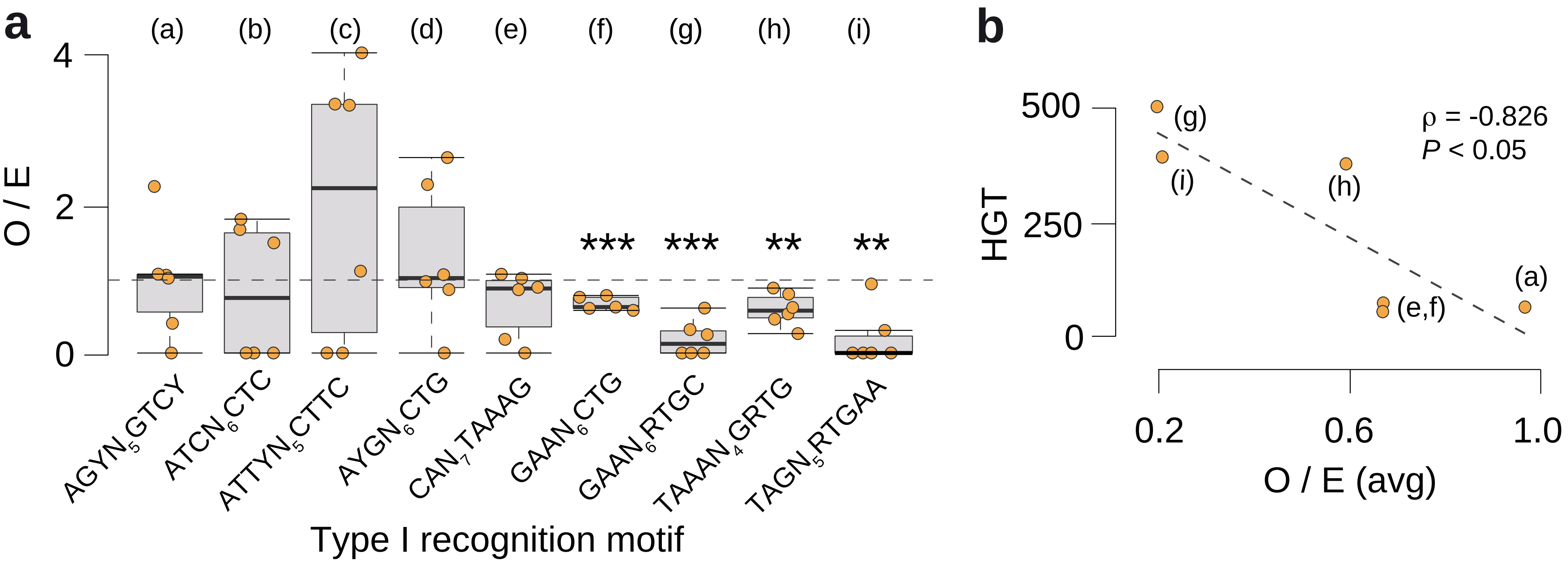

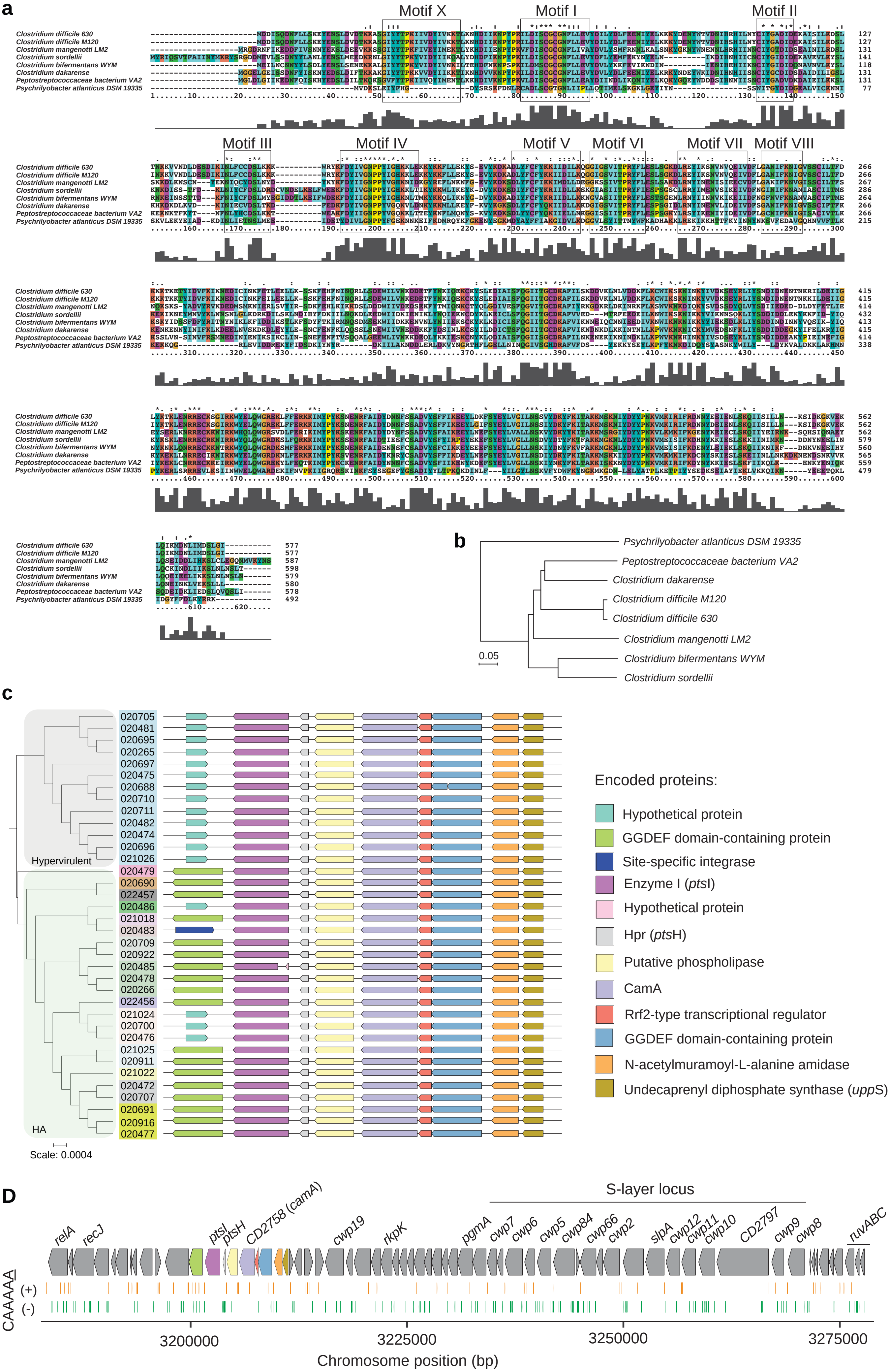

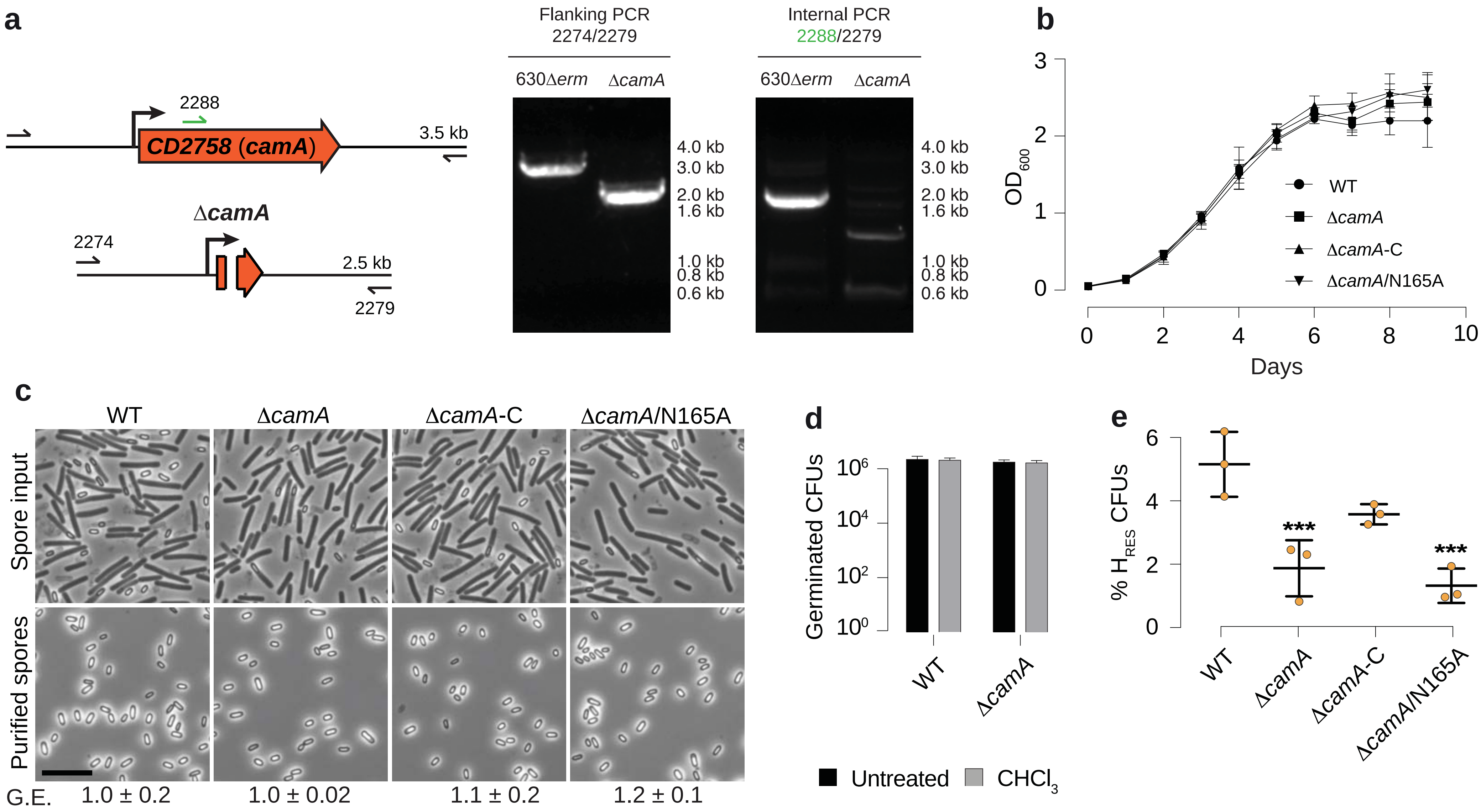

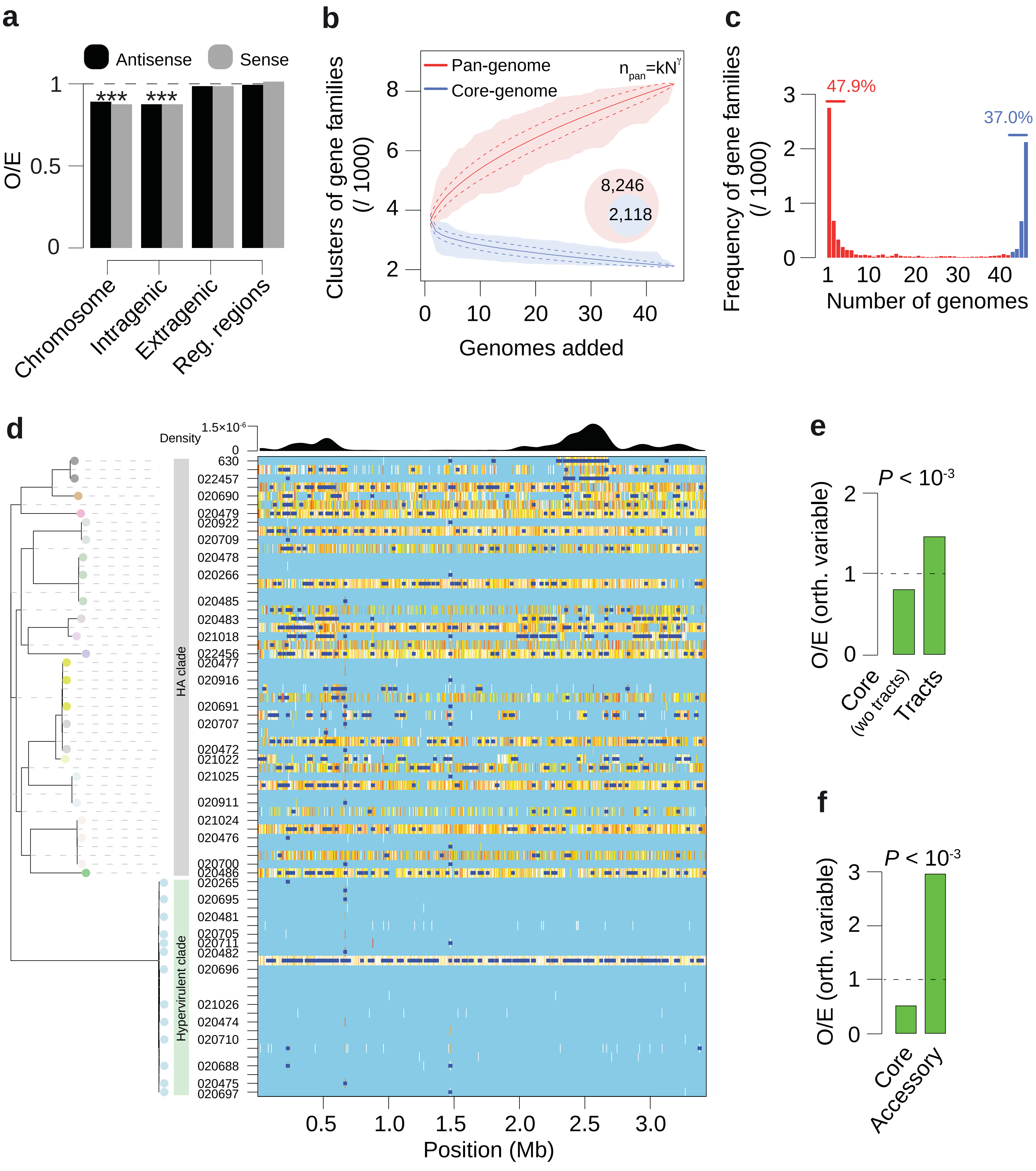

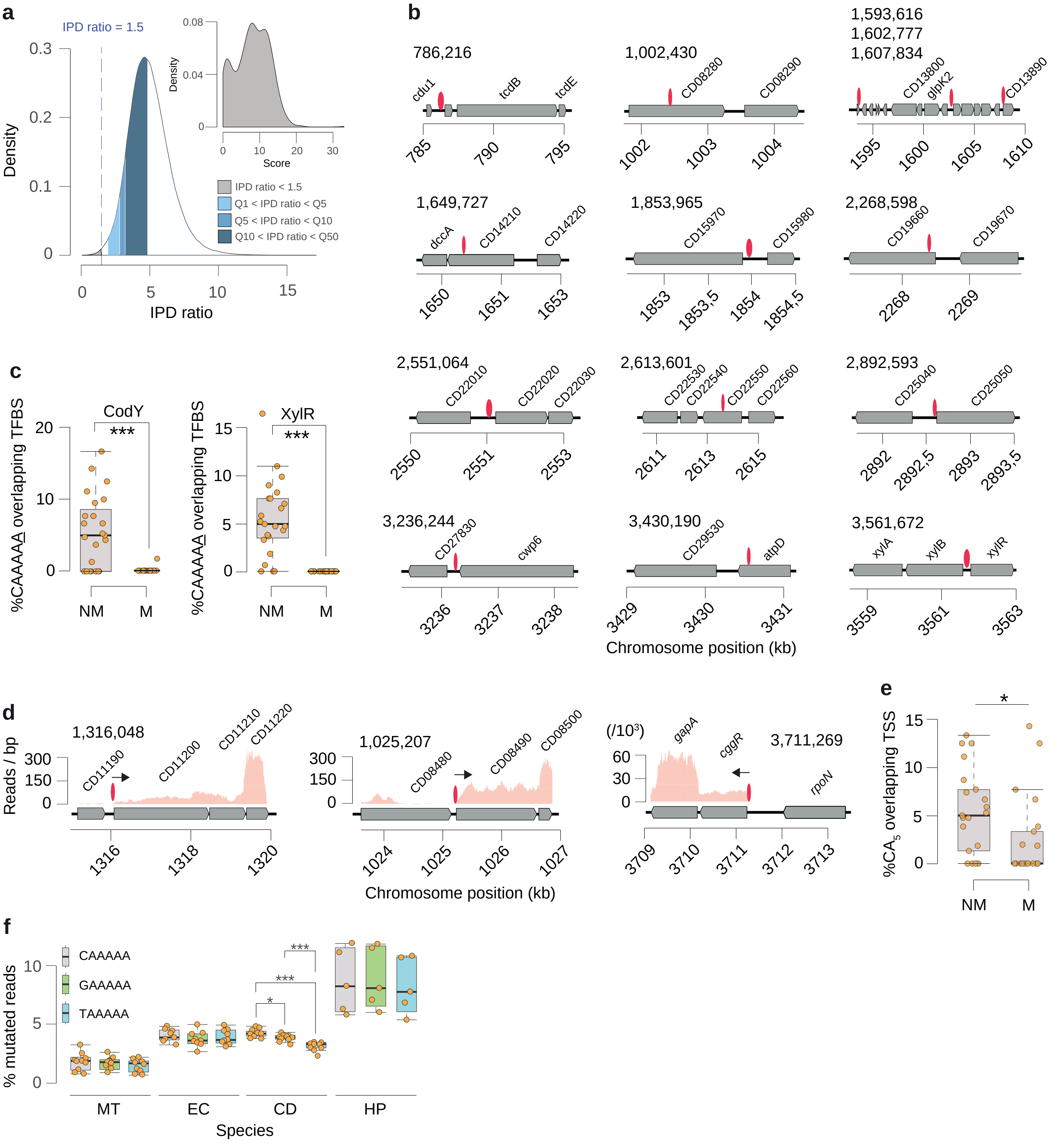

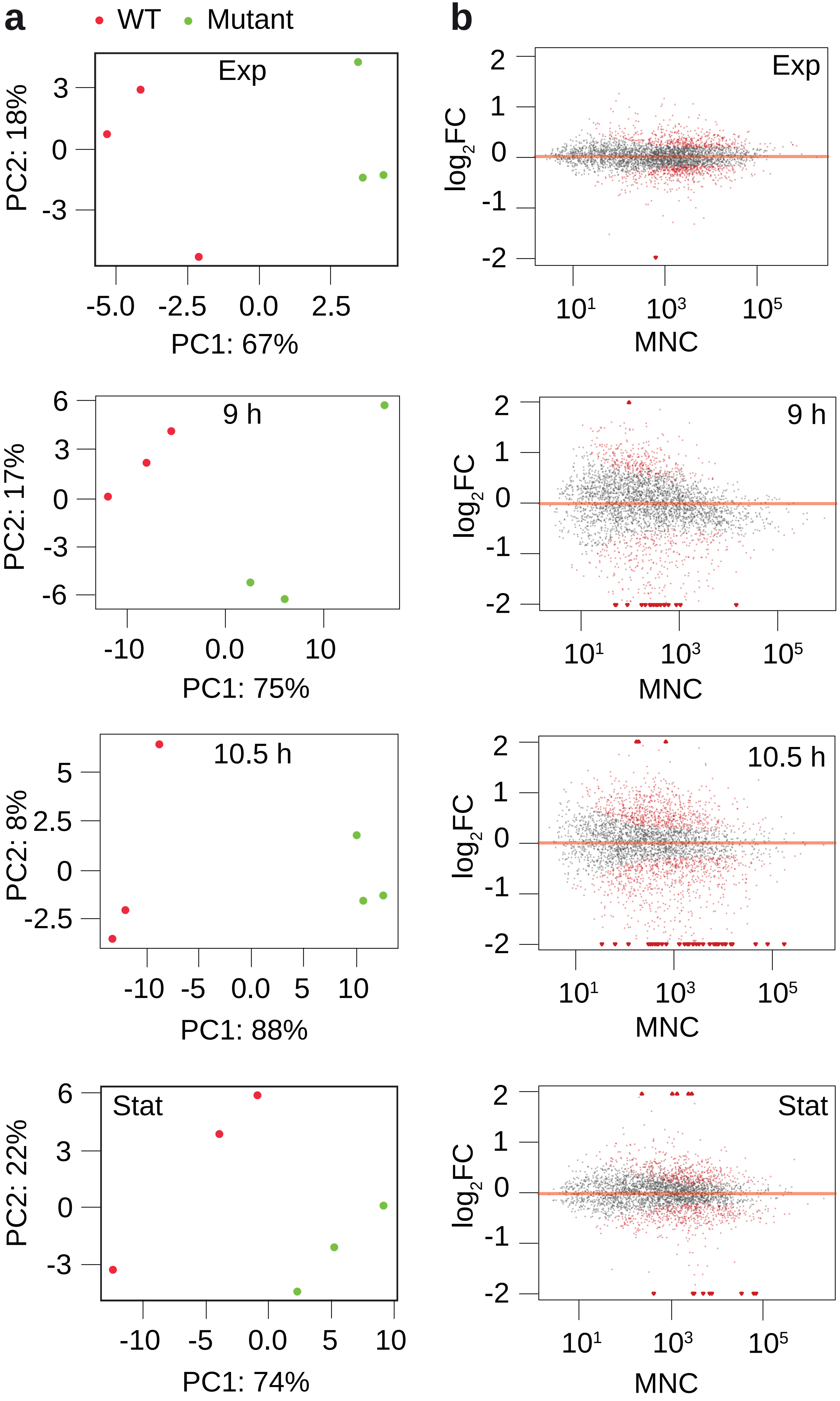

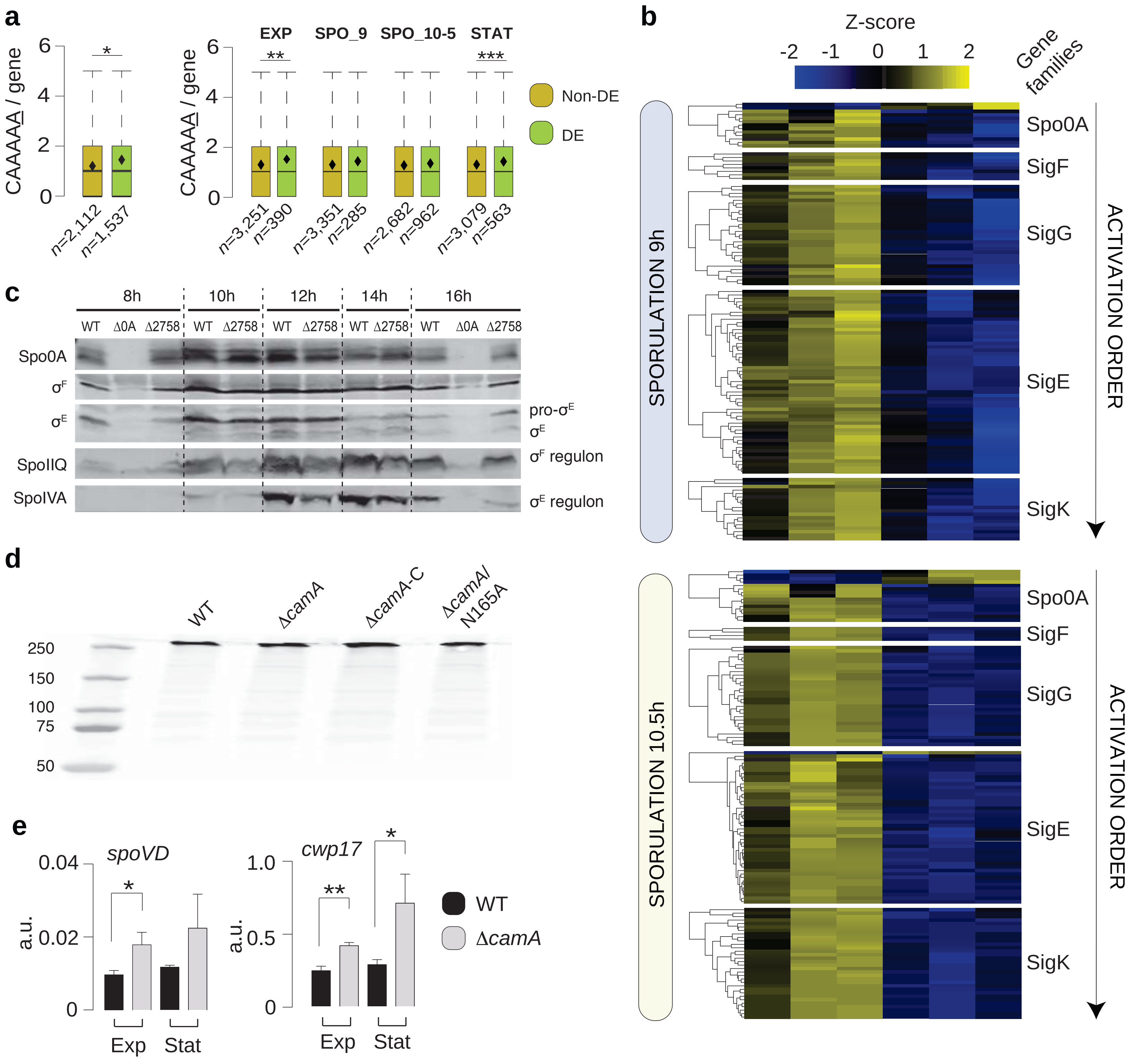

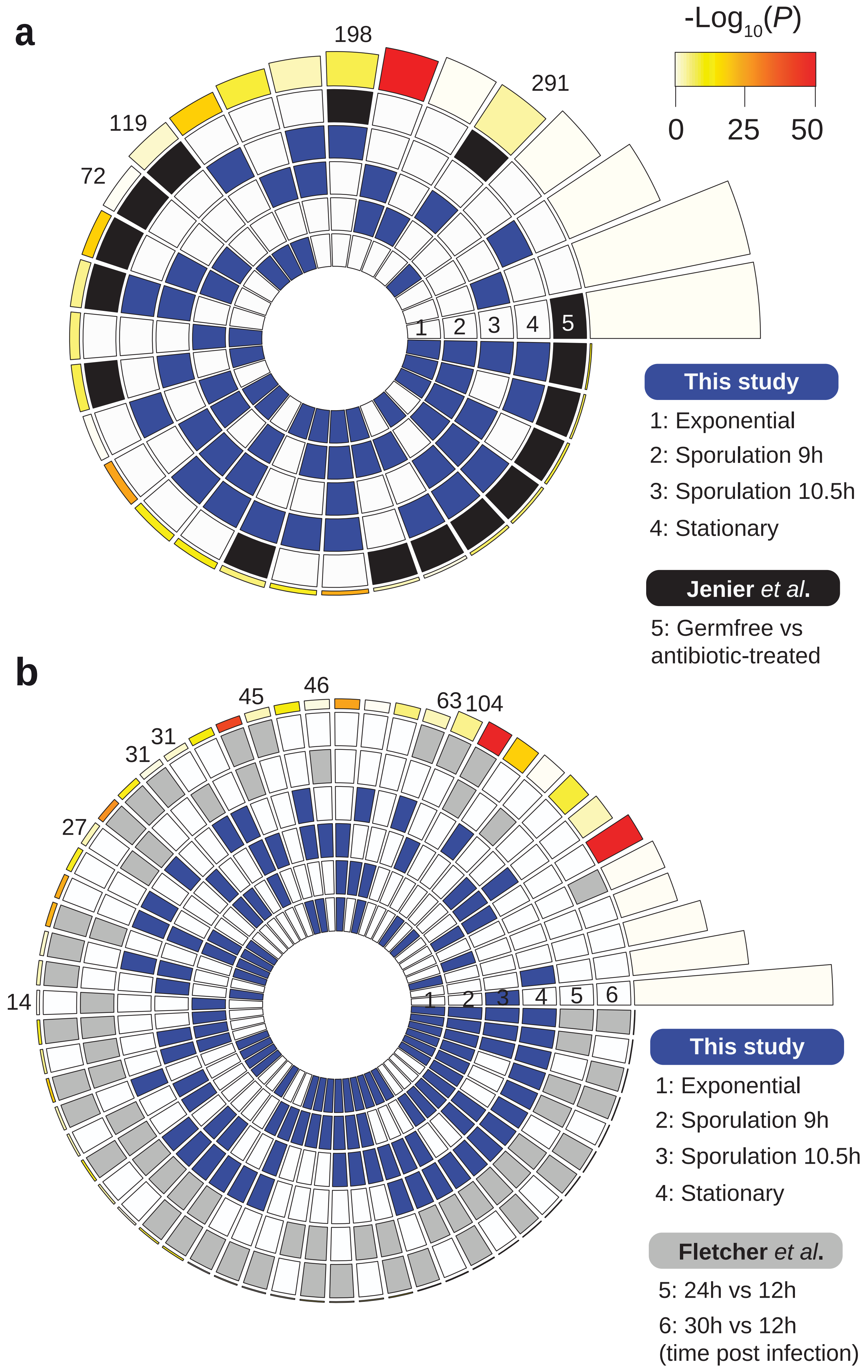
